# An Integrative Multi-Omics Random Forest Framework for Robust Biomarker Discovery

**DOI:** 10.1101/2025.03.05.641533

**Authors:** Wei Zhang, Hanchen Huang, Lily Wang, Brian D. Lehmann, Steven X. Chen

## Abstract

High-throughput technologies now produce a wide array of omics data, from genomic and transcriptomic profiles to epigenomic and proteomic measurements. Integrating these diverse data types can yield deeper insights into the biological mechanisms driving complex traits and diseases. Yet, extracting key shared biomarkers from multiple data layers remains a major challenge. We present a multivariate random forest (MRF)–based framework enhanced by a novel inverse minimal depth (IMD) metric for integrative variable selection. By assigning response variables to tree nodes and employing IMD to rank predictors, our approach efficiently identifies essential features across different omics types, even when confronted with high-dimensionality and noise. Through extensive simulations and analyses of multi-omics datasets from The Cancer Genome Atlas, we demonstrate that our method outperforms established integrative techniques in uncovering biologically meaningful biomarkers and pathways. Our findings show that selected biomarkers not only correlate with known regulatory and signaling networks but can also stratify patient subgroups with distinct clinical outcomes. The method’s scalable, interpretable, and user-friendly implementation ensures broad applicability to a range of research questions. This MRF-based framework advances robust biomarker discovery and integrative multi-omics analyses, accelerating the translation of complex molecular data into tangible biological and clinical insights.

## INTRODUCTION

Recent technological advances in high-throughput sequencing, mass spectrometry, and imaging have led to a surge in multi-omics data that span the genome, epigenome, transcriptome, proteome, and metabolome. Integrating these diverse data sources can provide a more comprehensive picture of complex biological systems than analyzing any single omics layer alone. When done effectively, integration can highlight shared molecular features across different data types, offering new insights into disease mechanisms, patient stratification, and potential biomarkers for clinical applications^1–4^.

Despite the promise of multi-omics integration, the process remains challenging. Traditional methods, such as sparse partial least squares (sPLS)^5,6^ and canonical correlation analysis (CCA) ^7–9^, focus largely on linear relationships. Although widely used, these approaches can struggle in high-dimensional settings, are prone to overfitting, and may fail to capture nonlinear interactions. Nonlinear extensions, including kernel CCA^10,11^, help address some of these issues but often face scalability and interpretability limitations, making them less suitable for many practical scenarios.

Ensemble learning techniques, particularly random forests, are valued for their robustness, ability to model nonlinearities, and relative resilience to overfitting^12^. Extending random forests to handle multiple response variables leads to multivariate random forests (MRF)^13^, which are well-positioned to tackle complex multi-omics data. However, applications of MRF to multi-omics integration have been limited, leaving an opportunity to develop methods that exploit the strengths of this approach for biomarker discovery and feature selection.

In this study, we introduce a new MRF-based framework that employs the inverse minimal depth (IMD) metric for variable selection across multiple omics datasets. By assigning response variables to decision tree nodes and using IMD to quantify feature importance, our method identifies key variables shared across different data layers. This strategy naturally reduces the risk of selecting noise variables and helps focus on those with consistent impact across datasets. Through simulations and applications to real-world datasets from The Cancer Genome Atlas (TCGA), we compare our method to established approaches like sPLS and CCA, demonstrating improved performance, stability, and interpretability. The biomarkers identified by our method correspond to meaningful biological pathways and can help distinguish patient groups with different outcomes.

In summary, our MRF-IMD framework provides a robust and flexible solution for multi-omics integration. By embracing nonlinear relationships, addressing high-dimensionality, and maintaining interpretability, this approach has the potential to advance biomarker discovery and contribute valuable insights to complex biological and clinical problems.

## MATERIAL AND METHODS

### Maximal Spitting Response Variable

Consider two datasets **X**_*n* ×*p*_ and **Y**_*n* ×*q*_ where *n* is the number of samples and *p* and *q* represent the number of features of ***X*** and ***Y*** respectively. Our goal is to integrate these datasets using a multivariate random forest (MRF) approach. In this framework, we use a splitting rule that considers all response variables together, rather than handling them individually. We begin with a splitting rule introduced by Tang and Ishwaran that extends the traditional univariate splitting criterion to a multivariate setting^18^. This rule extends univariate splitting by summing the splitting criterion across all response outcomes *Y*_.*j*_. The splitting criterion for node *t* is defined as:

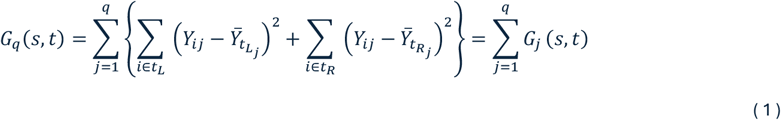

where 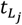 and 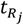 represents the left and right daughter nodes for *j*_*th*_ response coordinate and 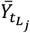 and 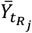 are the sample means in 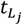 and 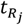. To determine the best split, we minimize *G*_*q*_(*s, t*), ensuring all response variables *Y*_.1_, …, *Y*_.*q*_ are measured on the same scale by standardizing them to a 0-1 scale: 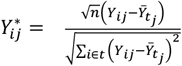, where

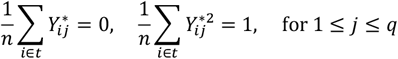

This standardization ensures that the contributions of all outcomes are comparable, preventing any single outcome from dominating the splitting process. After simplifying the expression, the minimization of *G*_*q*_ (*s, t*) becomes equivalent to maximizing:

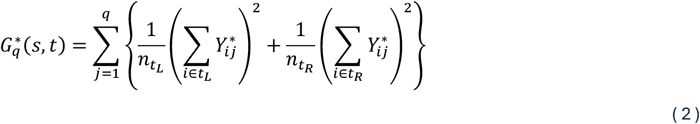

In the case of two-omics data, we treat one dataset as the response and the other as the predictor, and apply the multivariate random forest (MRF) model using these splitting rules.

Using the above framework, we now focus on two-omics data. For each response variable *Y*_*j*_, we define the splitting statistic as:

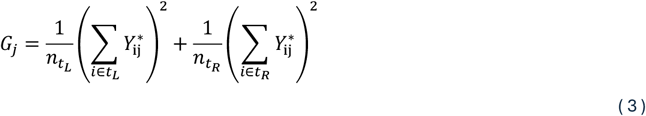

This statistic quantifies how well a split separates the values of *Y*_*j*_ in the response **Y** across the left and right daughter nodes. For each node split, we identify the maximal splitting response variable (MSRV) as the response variable that maximizes the multivariate splitting rule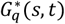, meaning it has the largest contribution to the split. The MSRV represents the variable most associated with the predictors at that particular node.

#### Definition 1

*In a multivariate tree T_b_ that has a total of m splitting, let* 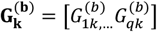 *be the splitting statistics in k_th_ splitting of the node, the MSRV* 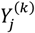 *in k*_*th*_ *splitting of a node is defined as the variable where* rgm 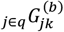, *among all variables in* **Y**.

#### Maximal Spitting Response Variable in Each Node Split

Once we identify the MSRV, we assign a single response variable from **Y** to each node in the tree. Intuitively, response variables that are more strongly associated with the predictors driving the node split will tend to have higher splitting statistics *G*_*j*_. In scenarios with sparsity, where only a few coordinates are non-zero, weak variables have a low probability of being selected as the MSRV—approximately 1/*q*, as selection in these cases can be largely random. To illustrate the MSRV, we used a random tree from an MRF model (Figure 1). The datasets X and Y were generated using the latent model, with the first 20 variables of each dataset being cross-correlated. The model was configured with *n* = 200 and *p* = *q* = 200. Nearly half of the MSRVs identified at each node are cross-correlated variables from Y, which are shown in yellow in the figure.

**Figure 1.**
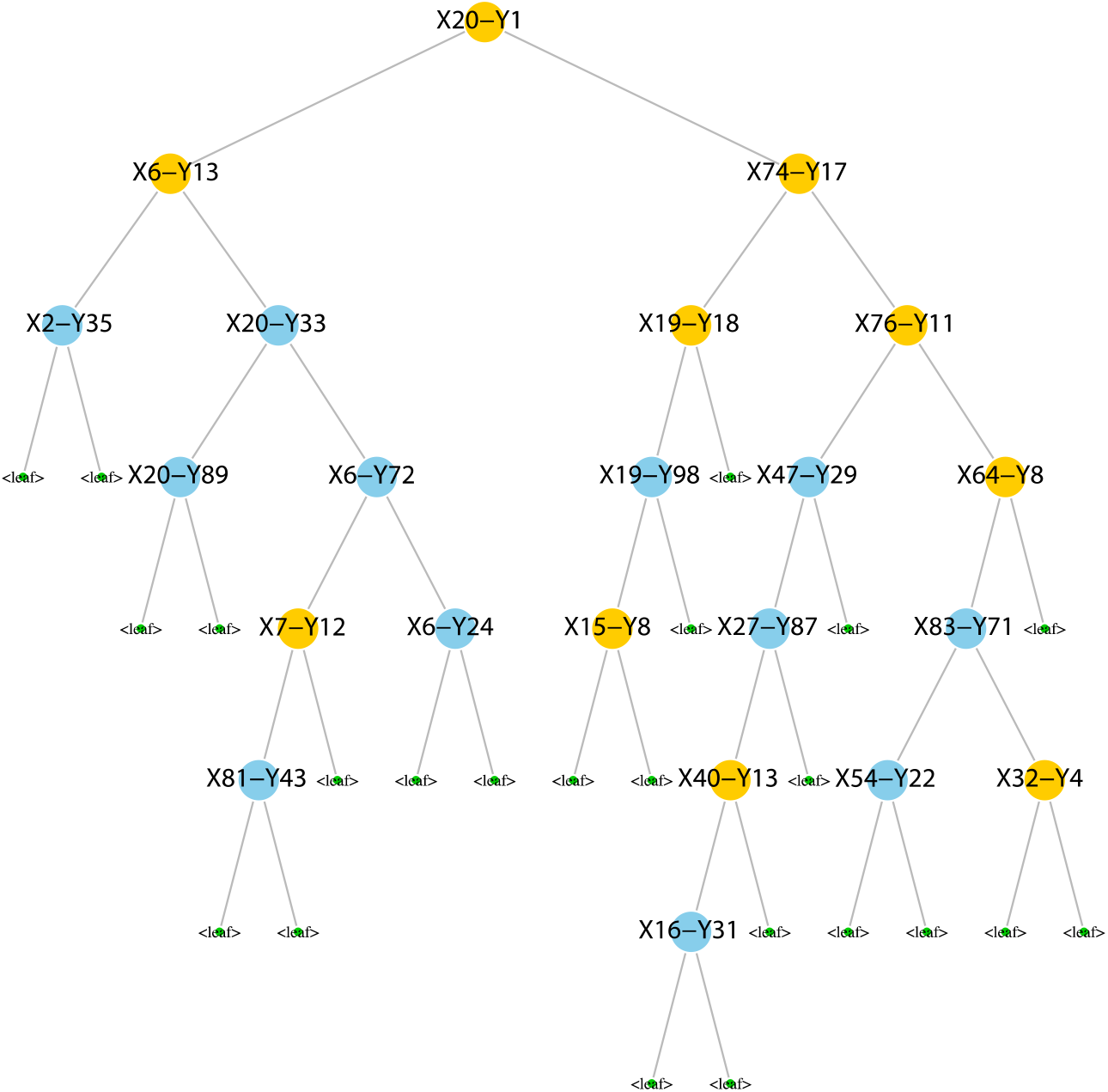
An illustration of a tree in MRF model. The datasets **X** and **Y** were generated using the latent model (Details in Simulation Study), with the first 20 variables of each dataset being cross-correlated. The variables *Y*_1_ to *Y*_20_ are cross-correlated with variables *X*_1_ to *X*_20_. Cross-correlated variables from **Y** are colored in yellow. Nearly half of the MSRVs identified at each node are cross-correlated variables from **Y**.

A detailed procedure for identifying MSRV is provided in the Algorithm 1 of Supplementary Note 1. In the high dimensional setting, calculating splitting statistics for all response variables can be computationally inefficient. Our primary goal in applying the MRF model to the two datasets is not prediction, but feature extraction to identify shared information. To address computational inefficiency, we select a random subset of response variables, which reduces complexity while still identifying key variables associated with both datasets. A key question is: what proportion of response variables should be selected at each node split? To explore this, we conducted two scenarios: (1) *q* = 200, where first 20 variables in both **X** and **Y** are cross-correlated, and (2) *q* = 500, where again the first 20 variables are cross-correlated. We set *p* = 500, *N* = 200, and *B* = 100 to control for other settings in both scenarios. For each proportion, we fit the MRF model and calculate the selection frequency with which variables in Y were selected as MSRV, repeating the procedure 50 times to ensure stability in the results.

Supplementary Figure S1 shows the box plot of the selection frequency across repeated trials, with cross-correlated variables highlighted in red. The plots indicate that, in both scenarios, cross-correlated variables are selected as MSRV more frequently than noise variables. On average, the selection frequency of noise variables **Y** as MSRV in a node is approximately 1/*q*, mirroring the probability of a noise variable in **X** being selected for a node split in sparse setting, which is around 1/*p*. The dashed line in the plots marked the 1/*q*threshold, and most noise variables cluster near this line. In Scenario 1, for proportions below 0.5, a few cross-correlated variables fall below the dashed line, while in Scenario 2, only the 0.1 proportion shows slightly more variables below the line. Scenario 1 also displays more variability in the selection frequency of cross-correlated and noise variables, suggesting that selecting MSRV is more reliable in sparse settings.

### Inverse Minimal Depth

#### Minimal Depth

Minimal depth, introduced by Ishwaran et al.^17,19^, is a variable selection method that efficiently ranks strong variables higher than weak ones. The minimal depth of a variable refers to the shortest distance from the root of a decision tree to the node where the variable appears. Let *D*_*v*_ denote the minimal depth of a variable *v* and *D*(*T*) represents the depth of a tree *T*, it has been proved that the distribution of *D*_*v*_ is :

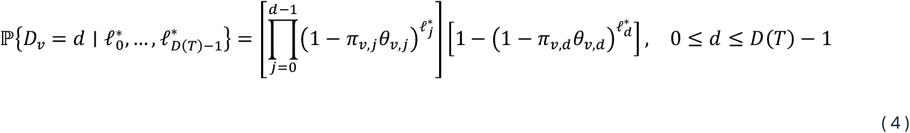

where *π*_*v,j*_ is the probability of *v* selected as a candidate variable for splitting of node *t* at depth *j, θ*_*v,j*_ is the probability of *v* splits a node *t* at depth *j*, and 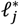 is the number of nodes at depth *j*. Note that if the tree is a balanced tree, 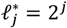 at depth *j*. For example, in Figure 1, the root node variable *X*_20_ is assigned a minimal depth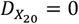. The left daughter node of *X*_20_, *X*_6_ has a minimal depth of 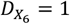 1. In the original paper on minimal depth, proposed two strategies for identifying strong variables. The first strategy uses the mean minimal depth under the null hypothesis that a variable *v* is a weak variable. Given *v* is a weak variable, the distribution of the minimal depth of *v* is

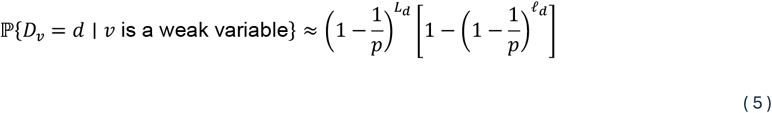

where *L*_*d*_ 1 + 2 + … + 2^*d*−1^ = ℓ_*d*_ − 1 and *p* is the number of features in the dataset. The threshold works well when *n* is large and is more computationally efficient than VIMP and jointly VIMP in high-dimensional MRF models. However, when dimensionality is high, meaning *p*≫ ℓ_*D*(*T*)_, the threshold will fail because all the probabilities ℙ{*D*_*v*_ *= d* | *v* is a weak variable} will approach 0. Later, we will demonstrate that the threshold fails in high dimensional noise settings with multivariate outcomes. In cases where the mean threshold approach fails, a second strategy known as variable hunting^19^. This approach involves iteratively selecting random subsets of variables, fitting the forest, and combining minimal depth with joint VIMP to prioritize the strongest variables. While effective when the number of features *p* greatly exceeds the number of samples (i.e. *p*≫ *n*), this method becomes computationally inefficient and can struggle in high-dimensional noise settings with multivariate outcomes. To overcome these limitations, we propose a variation of the minimal depth approach, which we outline in the next section. This variation allows for the selection of strong variables in both the response and predictor spaces within the multivariate random forest (MRF) model.

#### Distribution of Inverse Minimal Depth

To incorporate minimal depth into our variable selection method, we introduce a new statistic called inverse minimal depth (IMD), defined as:

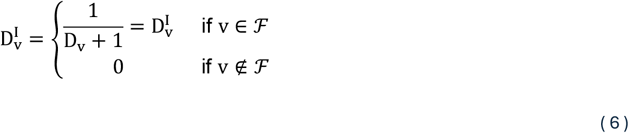

where ℱ is the set of variables selected in the tree. For example, as shown Figure 1, *X*_20_ has a minimal depth of 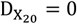 and its IMD is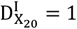. Similarly, *X*_6_ has a minimal depth of 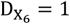 and its IMD is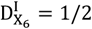. In this setup, larger IMD values correspond to stronger variables, making it easier to identify them. Additionally, we apply a penalization technique that assigns an IMD of 0 to variables that are not selected in the tree. When V ∈ ℱ, the distribution of D_V_ can be directly transformed from the distribution of as D_V_ follows:

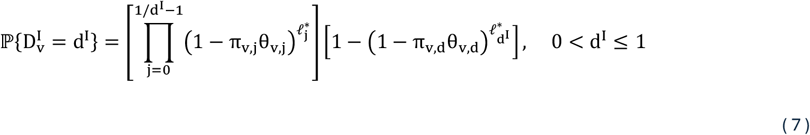

Note that 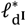 is the same value as in the distribution of minimal depth. With IMD, values are confined between 0 and 1. Variables not selected (V ∉ ℱ) have 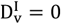, while stronger variables exhibit higher IMD values. The overall IMD for a variable V across the forest is calculated as the average IMD across all trees:

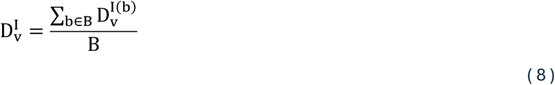

To further investigate the relationship between IMD and key MRF parameters—such as the number of trees, tree depth, and the proportion of response variables in each split—we conducted extensive simulations, detailed in Supplementary Note 2. Additionally, to compare the performance of the original MD and the enhanced IMD metric, we performed simulations outlined in Supplementary Note 3.

#### Variable Selection in High Dimensional Two-omics Data

As previously noted, one method for variable selection based on minimal depth involves using a pre-defined threshold derived from the distribution of weak variables’ minimal depth. However, when *p*≫ ℓ_*D*(*T*)_, all the probabilities in (5) approach zero^17^. Applying the same thresholding method to IMD yields similar challenges in high-dimensional datasets, where thresholding values also tend toward zero. To address this, we propose two additional methods for detecting strong variables that are not based on weak variable distributions. First, we introduce these approaches for two-omics data in a multivariate random forest. Then, we extend the selection framework to multi-omics variable selection.

#### Variable Filtering

As the IMD of noise variables are close to or hovers over 0, we can select the threshold by multiplying a parameter *τ* to the standard deviation of IMD: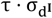. To determine the optimal value of τ, we use the mean out-of-bag errors (OOB) of both response and predictor variables for tuning. The mean OOB error is averaged across all the OOB errors in **X** and **Y**. In the MRF setting, for each i_*th*_ response in **Y**_n×q_, the loss function becomes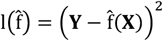. Therefore, the estimation of prediction error for response the OOB sample can be defined as:

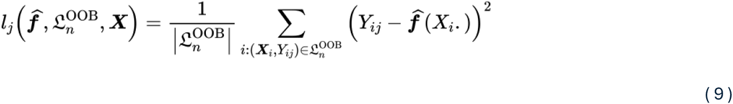

Similarly, the OOB errors of predictors **X** can be derived from the forest weights statistics. The OOB prediction of **X** can be formulated as follows:

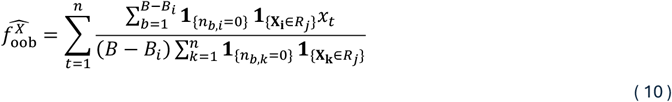

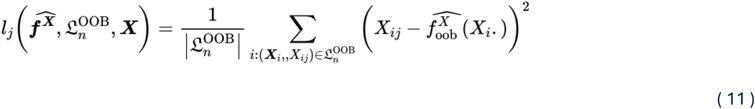

With a step size of 0.1, we computed the mean OOB error using the variables above 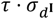. To stabilize the OOB error, we repeated the model fittings *k* times and averaged the results to select the optimal *τ* based on a tolerable error deviation.

#### Detecting Signals with Mixture Model

Given the distribution of differences in IMD between strong variables and noise variables, we can identify strong cross-correlated variables by fitting a two-component mixture model to the forest IMD. We describe the univariate Gaussian mixture model as follows:

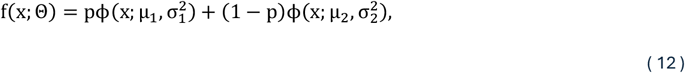

where ϕ(·) is the normal distribution. As the forest IMD ranges from 0 to 1, we consider modeling the IMD using truncated distribution. A previous study used the truncated normal mixture model to model the intraclass correlation coefficient of DNA methylation probes. The distribution of the truncated normal mixture model is as follows:

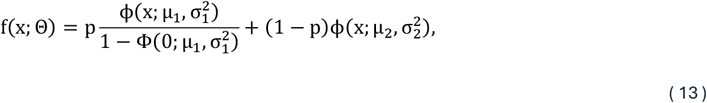

where *p*∈ [0,1] is the proportion of the first component, and the intraclass correlation is bounded by (0,1). Here, we assume that the noise variables have relatively low forest IMD and more likely lie in the first component modeled by the normal or truncated normal distribution. To estimate the parameter of the mixture model, we used the Expectation-Maximization (EM) algorithm for the model fitting^20,21^. To accommodate the variables with forest IMD = 0, we used the modified log-likelihood function proposed in:

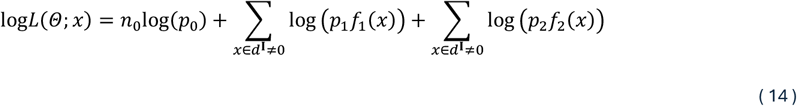

where *p*_0_ is the proportion of forest IMD = 0, *p*_1_ = *p*(1 − *p*_0_), and *p*2 = (1 − *p*)(1 − *p*_0_). In the forest IMD, we separated the forest IMD = 0 and modeled the forest IMD > 0 using (12) or (13). Figure 2 shows the density of modeling the forest IMD using Gaussian mixture and truncated normal mixture models. The data is simulated by the latent model with the settings of *p* = *q* = 500, *n* = 200, and the first 30 variables of each dataset are cross-correlated with each other. Here, we can see that the forest IMD has a skewed distribution. The Gaussian mixture model shows a better fit of the forest IMD. For each component, the posterior probabilities can be calculated as:

**Figure 2.**
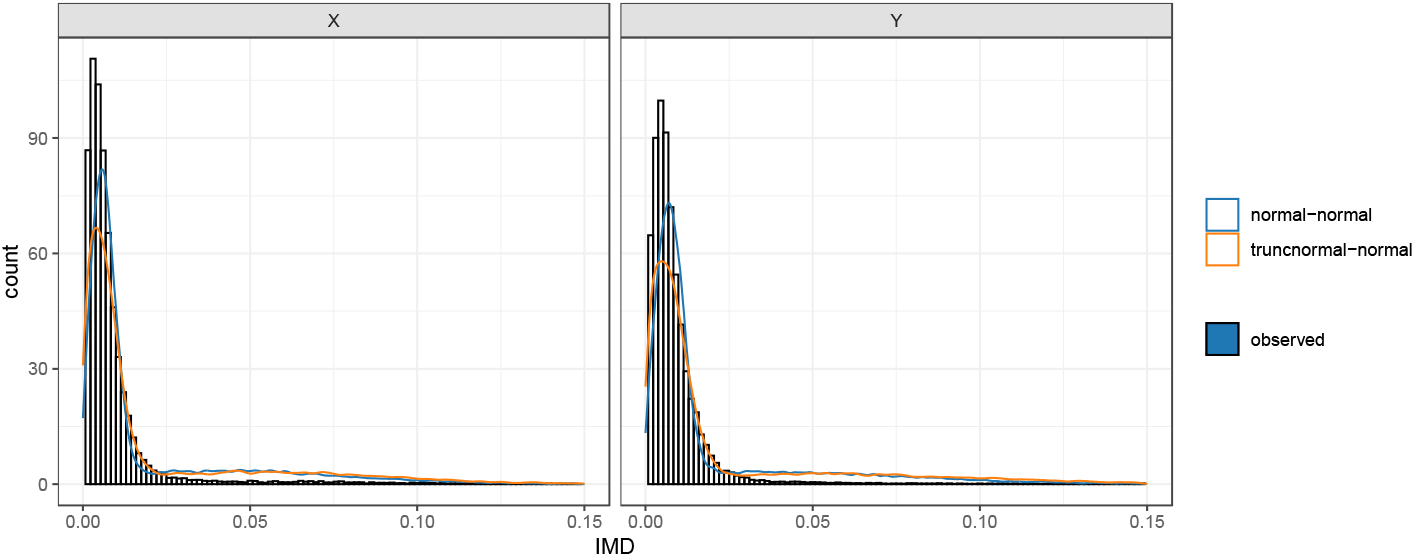
Modeling forest IMD with Gaussian mixture (blue curve) and truncated normal mixture (orange curve) models. The data is simulated by the latent model with the settings of *p* = *q* = 500, *n* = 200, and the first 30 variables of each dataset are cross-correlated with each other. The forest IMD has a skewed distribution, and the Gaussian mixture model fits better than the truncated normal mixture in this scenario.

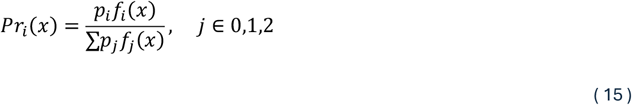

We selected variables with *pr*_1_(*x*) < *pr* as the important variables, where *pr* is a predefined value (i.e., *pr* = 0.05).

#### IMD Transformation

In the previous section, we discovered that the mean IMD of strong variables is close to the mean IMD of noise variables in high-dimensional settings. However, the forest IMD of strong or cross-correlated variables may be low and mingled with noise variables. This can be challenging for the mixture model to capture due to the sparsity of strong variables. To address this issue, we propose a third method that explores the distribution of IMD for both noise and strong variables. Using the latent model, we simulated two datasets with the same settings as in the previous section. From Figure 3a, it is clear that the forest IMD of noise variables skews heavily toward the lower end of the IMD scale, clustering near zero. In contrast, the cross-correlated variables exhibit a broader distribution starting from zero, although this is less apparent due to their sparsity. This makes distinguishing between noise and strong variables using only a threshold on the original forest IMD difficult. Let *µ* (displayed as a black dashed line), *µ*_*Noise*_ (displayed as a red dashed line), and *µ*_*Strong*_ (displayed as a blue dashed line) denote the mean forest IMD of all variables, noise variables, and cross-correlated variables, respectively. It is clear to see that *µ*_*Noise*_ ≤ *µ* ≤ *µ*_*Strong*_ and that *µ*_*Noise*_ tends towards *µ*.

**Figure 3.**
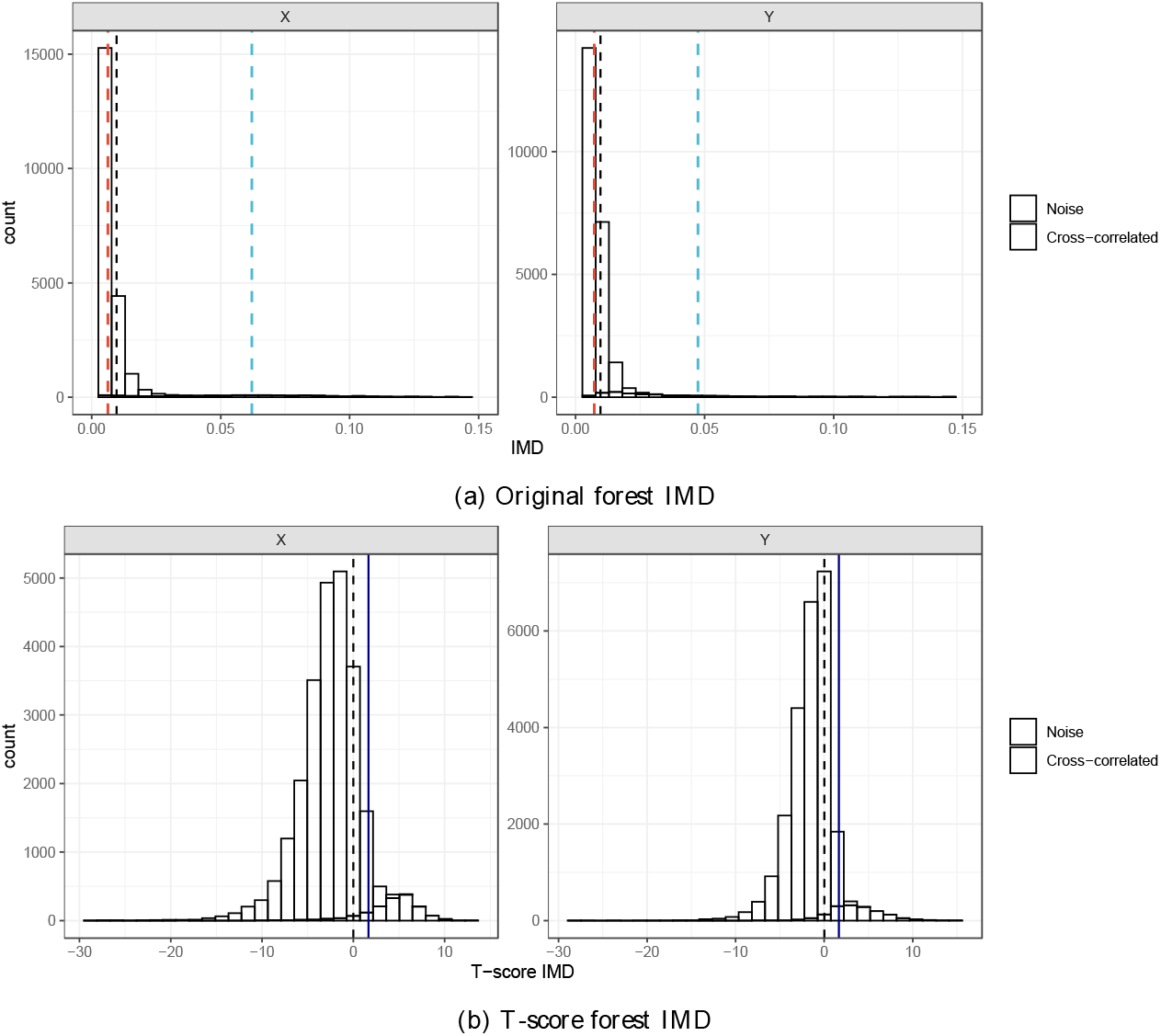
Distribution of IMD of cross-correlated and noise variables. The (a) original forest IMD of noise variables skews heavily toward the lower end of the IMD scale, clustering near zero. In contrast, the cross-correlated variables exhibit a broader distribution starting from zero, although this is less apparent due to their sparsity. The (b) t-score forest IMD effectively differentiates between noise and cross-correlated variables. Noise variables cluster below the mean (μ), while cross-correlated variables significantly diverge from μ.

Based on these findings, we standardize the forest IMD of variable *v* using the mean *µ* and the standard error of IMD of *v*. The standardization is represented by:

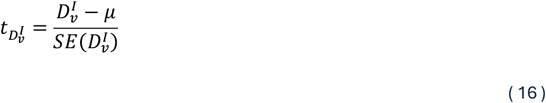

We denote 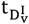 as the t-score IMD of variable v. This transformation yields a symmetric distribution (Figure 3b). The t-score IMD effectively differentiates between noise and cross-correlated variables. Noise variables cluster below the mean (μ), while cross-correlated variables significantly diverge from μ. Using the lower tail of the t-distribution at the 0.05 level (t_0.05,df=ntree−1_, denoted by the navy line), we identified the cross-correlated variables that have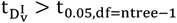. This specific threshold represents a point below which only an expected 5% of the IMD values for cross-correlated variables fall, thus indicating a higher level of importance.

#### Multi-omics Framework

We now extend the variable selection framework to multi-omics data. While we have introduced the variable selection method in two-omics data, it is essential to recognize that the choice of which dataset to assign as responses or predictors can influence the results, particularly in complex datasets. Additionally, the strength of connections between datasets plays a critical role in multi-omics variable selection, as datasets may share little common information. In such cases, variable selection can become biased. This section introduces an algorithm designed to efficiently find optimal connections between multi-omics datasets. To improve computational efficiency and address the high-dimensionality challenge in multi-omics data analysis, we applied principal component analysis (PCA) to each dataset. PCA reduces the dimensionality of the datasets by selecting components that explain a predefined level of cumulative variance. This ensures that we retain the most relevant information while minimizing computational complexity. For each reduced dataset, we conducted multivariate random forest (MRF) modeling, matching each dataset as a response to all others as predictors. We evaluated these models by calculating the mean out-of-bag (OOB) error, which was used to rank the models. The direction with the lowest OOB error was selected as the optimal connection between the datasets. This process enhances efficiency by avoiding exhaustive pairwise modeling and retains the most informative variables for further analysis.

For multi-omics datasets, let χ {**X**^(1)^, **X**^(2)^,…, **X**^(*K*)^} denote a multi-omics dataset with *K* omics data, where 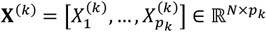 denotes the *k*_th_ omics data with *N* data samples and *p*_k_ features. Let ℳ be the the model collection that contains all optimal connected MRF models and 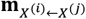 is the model in ℳ with the direction of *X*^(*i*)^ as responses and *X*^(*j*)^ as predictors. Algorithm 2 in Supplementary Note 4 summarized the framework of multi-omics variable selection under the variable filtering and mixture model methods. First, we compute the mean IMD for each dataset across model set ℳ. Instead of individual IMD, we select the important variables based on the mean IMD. For multi-omics variable selection under the IMD transformation, we choose the variables that the majority of the model selects (See Algorithm 3 in Supplementary Note 5).

#### Simulation Study

To evaluate the performance of our proposed methods in variable selection, we conducted a comprehensive simulation study. We generated synthetic multi-omics datasets under various conditions to assess the impact of noise, high dimensionality, and different proportions of cross-correlated variables on the accuracy of our model. The simulation was designed to mimic realistic data integration challenges, where datasets contain a mixture of relevant and irrelevant variables. We compared our IMD-based methods with existing techniques such as sparse PLS (SPLS), penalized matrix decomposition CCA (PMDCCA), and sparse regularized generalized CCA (SGCCA), assessing their ability to select important variables across different scenarios.

We designed a comprehensive simulation study to evaluate the performance of the proposed IMD-based methods under various conditions. Two different models were used: a **latent model** and a **non-linear regression model**. For each model, we generated synthetic multi-omics datasets to assess the impact of dimensionality, noise, and varying proportions of cross-correlated variables on model performance.

#### Latent Model

For the simulation of the linear models, we use the following model:

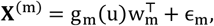

where 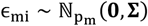 for i 1, …, p_m_, and u is generated from normal distribution u ∼ N(**0**, σ^2^) using mean 0 and σ = 2. For the kernel functions, we set g_1_(μ) = μ^2^, g_2_(μ) e p(μ), and g_3_(μ) = μ to transform the latent variable μ. The weights variable w_m_ were first generated by variables 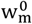 from uniform distribution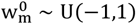.). Then, the variables were normalized in the following way to ensure that the sum of squared of w_m_ is equal to 1:

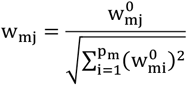

For this model, we generated two-dataset and three-dataset settings. For each dataset, only the first 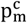 of weights w_m_ are non-zero and selected as the features to identify. Scenarios were generated based on the following parameters: 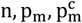, where n represents the sample size of all datasets, p_m_ represents the feature numbers of **X**^(m)^, and 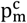 the number of features that crossed-correlated w.r.t. **X**^(m)^. Here we set the diagonal of the variance-covariance matrix **Σ** to 0.3^2^ and all variables in the **X**^(m)^ were re-scaled to have a mean of 0 and a standard deviation of 1. To generate data, we used the scenarios described in Table 1a for the two- and three-data settings.

**Table 1:** Simulation model scenarios.

(a) Latent model
| Scenario | S1 | S2 | S3 |
| --- | --- | --- | --- |
| Sample size | 100 | 200 | 200 |
| Dimension | 200 | 500 | 1000 |
| True model size |  |  |  |
| Two-omics | 20 | 30 | 50 |
| Three-omics |  |  |  |

(b) Non-linear regression model
| Scenario | S1 | S2 | S3 |
| --- | --- | --- | --- |
| Sample size | 100 | 200 | 200 |
| Dimension | 200 | 500 | 1000 |
| True model size |  |  |  |
| Setting 1 | 20/5 |  |  |
| Setting 2 | 40/10 |  |  |

#### Non-linear Regression Model

This simulation model was inspired by the simulation model in Degenhardt et al.^22^. Let be the basis variables that are generated from a distribution. For this model, we will add four additional parameters, 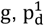, and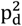, where g represents the group size of each correlated group in **X**, and 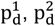 represent the number of variables that are not cross-correlated in **X** and **Y** respectively. This time, we use p to represent the total number of basis that formed **Y**. To generate the correlation between **X**, the simulation is according to:

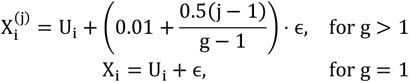

for i = 1, …, g and i = 1, …, p, where *X*^()^ denotes the _*th*_ variable in group i. Note that when g = 1, there is no correlation between *X*_i_. When g > 1, there will be g correlated variable in variable group i, the increase in will decrease the correlation between basis variable 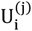 and *X*. And notice that the total number of features that crossed-correlated variables in 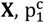, is equal to g · p_1_. To generate **Y**, we use the following kernel function:

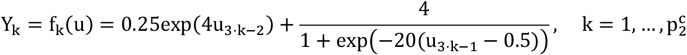

where each cross-correlated is formed by 2 basis variables for a total of 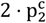 number of basis.

Therefore, 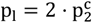.

Finally, we will generate two sets of noise variables for **X** and **Y** respectively using multivariate Gaussian distribution with mean 0 and identity variance-covariance matrix. These two sets of variables are neither cross-correlated nor inner-correlated to each other. The number of independent sets for **X** is 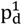 and the number of independent sets for **Y** is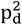. Note that all variables in the **X** and **Y** matrix are re-scaled to have mean 0 and standard deviation 1 after generating by the above models. We generated two settings of cross-correlated variables, and in each setting, we generated three different dimensional scenarios as shown in Table 1b.

### Evaluation Metrics

For each simulation scenario, we evaluated the performance of the variable selection methods using the following metrics: recall, precision, area under the precision-recall curve (PR-AUC), and model size.

#### Recall

This metric measures the proportion of true important variables identified by the model. Let *t*p be the number of variables the model selected as important that are true important variables, and fn be the number of variables that the model selected as important variables that are noise variables. The recall equals 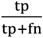. A higher recall indicates that more important variables were selected.

#### Precision

This metric measures the accuracy of the variable selection, focusing on the proportion of correctly identified important variables out of all selected variables. Let fp be the number of variables that the model selected as important that are true non-important variables, and *t*p described above. The precision equals 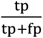. Higher precision indicates that fewer noise variables were incorrectly selected.

#### PR-AUC

This is a common metric for imbalanced datasets where important variables (positive cases) are fewer compared to noise variables (negative cases). PR-AUC ranges from 0 to 1, with a value of 1 indicating a perfect classifier. We reported the average PR-AUC across all datasets.

#### Model size

This represents the number of variables selected by the model. We calculated the average model size for each simulation scenario and reported the standard deviation to capture the variability in model size across replicates. For stability, each scenario was simulated 50 times, and all evaluation metrics (except model size) were averaged across replicates. For model size, we reported the average and standard deviation to assess the variability in the number of selected variables.

All MRF methods used the same parameter setting with and default settings from the rfsrc function in randomforestSRC R package throughout all simulation scenarios. All other methods have a tuning function for selecting the best number of variables in their packages. For SPLS method in mixOmics package, it contains a tuning function for selecting the best number of variables to keep. The PMA and RGCCA packages have tuning functions for selecting the penalty terms using a permutation test.

### TCGA data

To demonstrate the effectiveness of our proposed methods, we applied our methods to three TCGA datasets: breast invasive carcinoma (BRCA)^23^, colon adenocarcinoma (COAD)^24^, and Pan-Cancer^25^. For TCGA-BRCA and TCGA-COAD data, we analyzed three types of omics data (i.e., mRNA expression data (Gene), miRNA expression data (miRNA), and DNA methylation data (Methyl)) were selected to perform analysis. For TCGA-Pan-Cancer, we selected the ATAC sequencing data and RNA sequencing (RNA-seq) datasets. All the RNA-seq data, which were log2-transformed transcripts per million (TPM) for each cancer type, were obtained from the R package *TCGAbiolinks*. Other datasets were downloaded through UCSC Xena (https://xena.ucsc.edu/)^26^. For TCGA-Pan-Cancer, we selected the ATAC sequencing data and RNA sequencing datasets. Each of these data types provides unique, yet complementary, information for distinguishing between different types or states. Analyzing RNA-seq and ATAC-seq independently, however, can lead to inconsistent classifications. Furthermore, studying these two modalities in isolation may diminish the overall power of the analysis, as they both represent the same fundamental types or states. Only samples that existed across all data types were included in our study. We applied log2-transformation to mRNA and miRNA expression data. For DNA methylation, we initially filtered out probes not included in the Illumina Infinium HumanMethylation450k BeadChip to ensure a better interpretation of results. For miRNA, we removed all the variables that contain missing values. To streamline our analysis, we limited the features of mRNA and DNA methylation data to the top 2000 most variable expressions in both the BRCA and COAD datasets. For the Pan-Cancer dataset, we restricted ATAC-seq and RNA-seq to the top 50,000 and 5,000 most variable expressions, respectively. Table 3 summarizes the datasets used in the analysis.

## RESULTS

### Evaluation of simulated data

We first assessed the performance of our MRF-IMD framework on a variety of simulated scenarios designed to mimic common challenges in multi-omics integration. The simulations included both latent and nonlinear regression models, varying in sample size, dimensionality, and the proportion of truly cross-correlated variables between datasets. Our comparisons focused on three versions of our method—IMD-filter, IMD-mixture, and IMD-transformation—against established approaches such as sparse partial least squares (SPLS), penalized matrix decomposition CCA (PMDCCA), and sparse regularized generalized CCA (SGCCA).

Across the tested scenarios, all three IMD-based methods showed strong and stable performance. They consistently outperformed SPLS, PMDCCA, and SGCCA in terms of recall, precision, and PR-AUC, even as data dimensionality and noise levels increased. Notably, while reference methods performed acceptably under certain conditions, their accuracy and stability often declined as dimensional complexity grew. In contrast, MRF-IMD methods maintained reliable detection of cross-correlated variables, indicating their robustness to challenging data structures.

In simulations based on the latent model (Figure 4, Table 2), the IMD-mixture and IMD-transformation methods offered particularly balanced performance. They retained high precision and recall while achieving stable PR-AUC values over a wide range of parameter settings. Even though SPLS and RGCCA methods were developed under assumptions similar to those of the latent model, MRF-IMD methods remained competitive and often achieved better overall performance. This demonstrated that the IMD-based framework was not merely suited for one class of models, but could also adapt well to conditions where linear methods might be expected to excel.

**Table 2:**
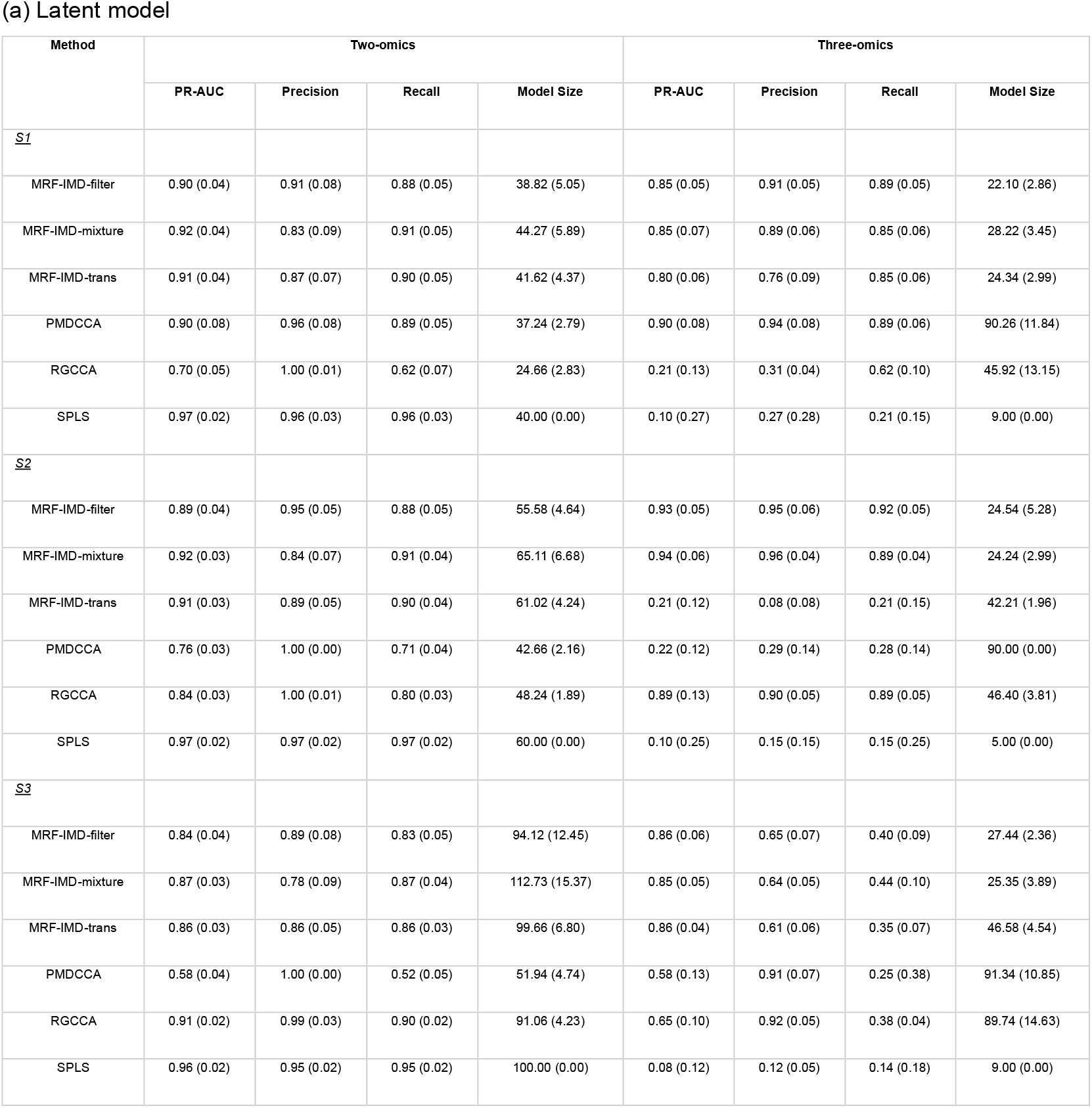
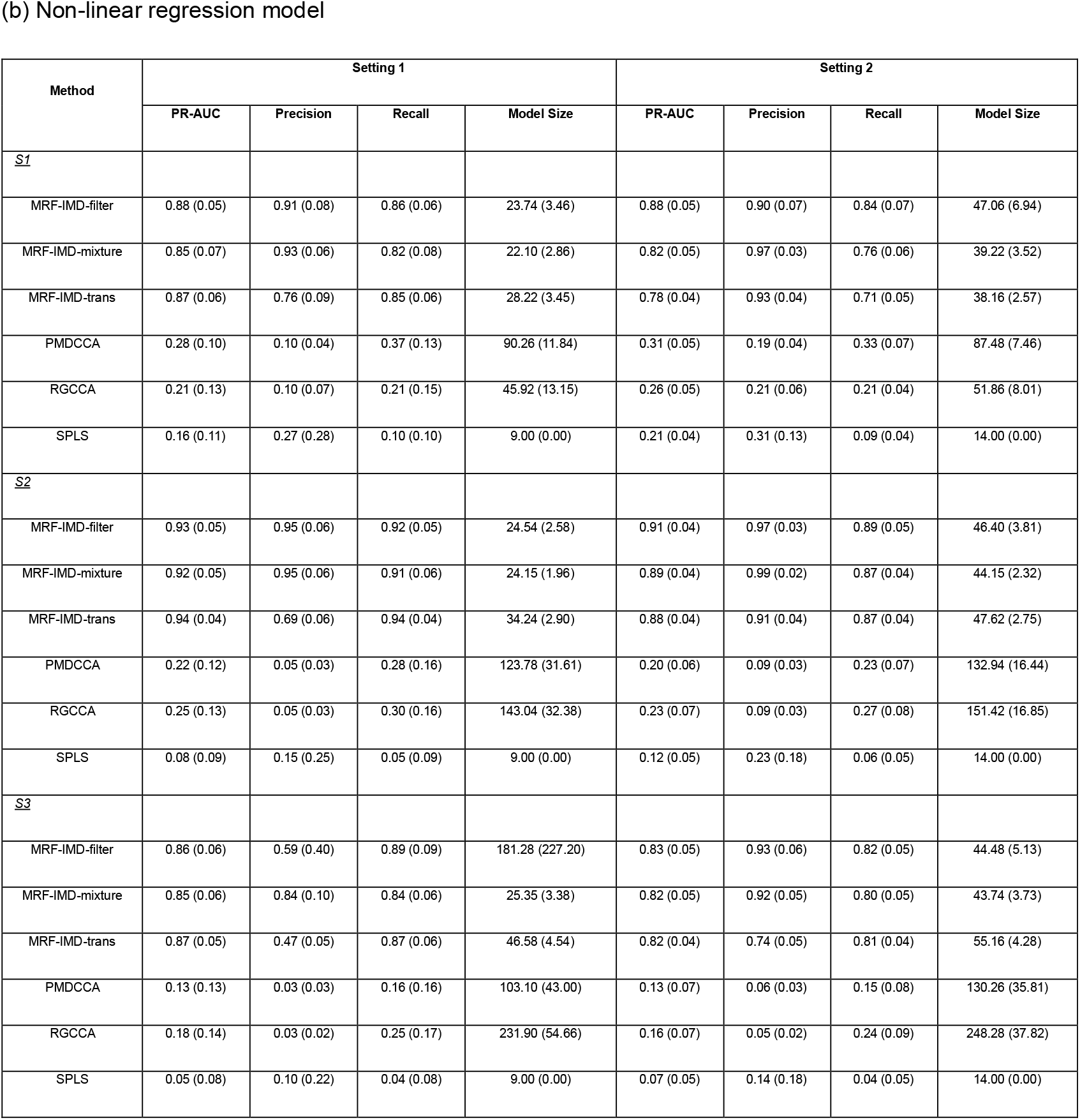
Simulation results. Performance measures are the mean of PR-AUC, precision, recall, and model size (standard deviations).

**Table 3:** Summary of datasets.

| Dataset | Number of arrays | Number of arrays for training | Number of samples |
| --- | --- | --- | --- |
|  | mRNA, DNAm, miRNA |  |  |
| BRCA | 20530, 485577, 2238 | 2000, 2000, 228 | 674 |
| COAD | 20530, 485577, 2113 | 2000, 2000, 252 | 257 |
|  | ATAC-Seq, RNA-Seq |  |  |
| Pan-Cancer | 562709, 59390 | 50000, 5000 | 383 |

**Figure 4.**
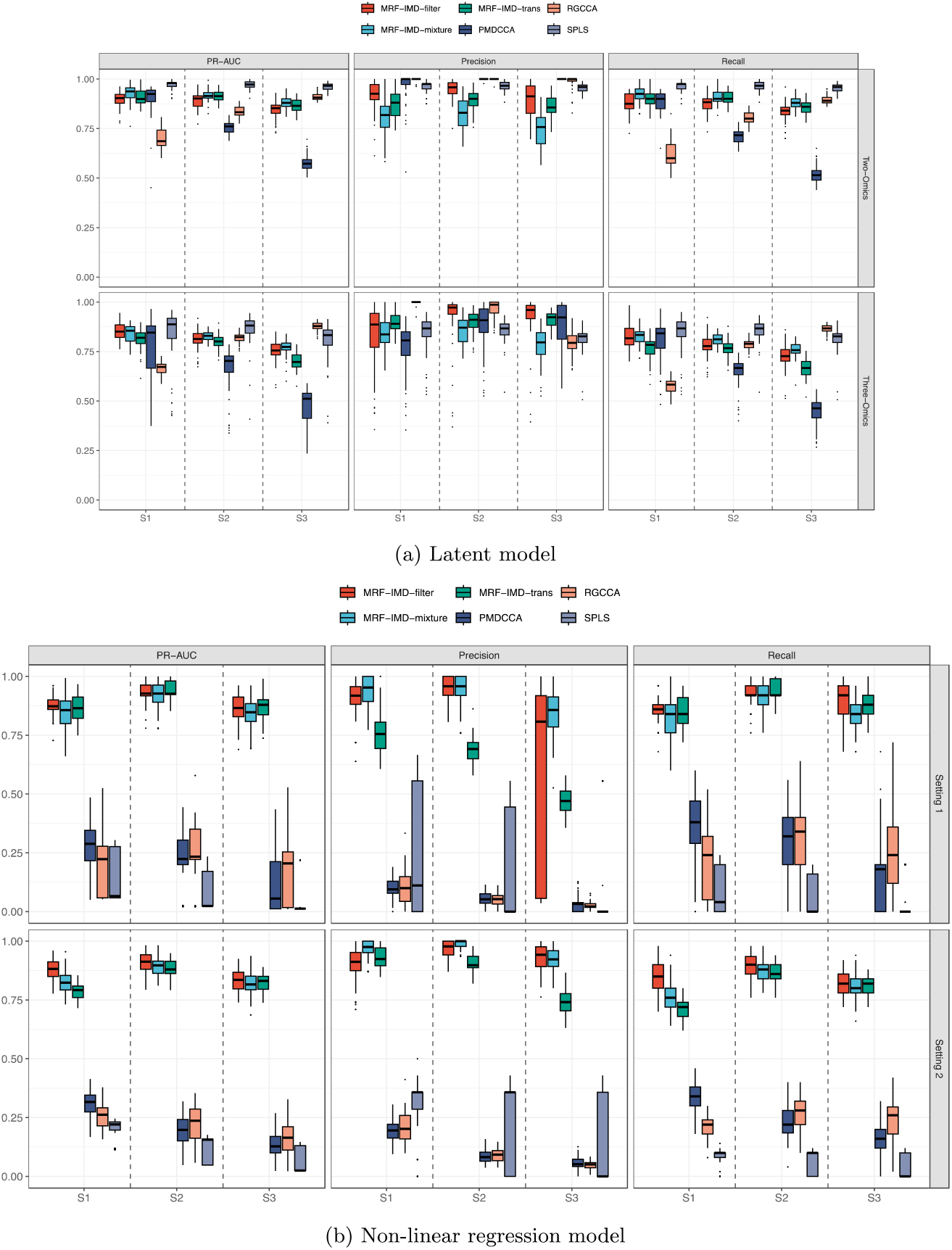
Boxplots of simulation results. Performance measures are PR-AUC, precision, and recall. In most scenarios, the MRF-IMD methods showed competitive selection outcomes in both (a) Latent model and (b) Non-linear regression model.

When tested on nonlinear regression models, the advantages of our MRF-based methods became more pronounced. Traditional linear methods struggled to capture the underlying nonlinear relationships, leading to reduced accuracy and higher variability. In contrast, the IMD-filter, IMD-mixture, and IMD-transformation approaches remained steady, offering consistently high PR-AUC values and stable variable selection results. These outcomes highlight the benefit of modeling complex interactions through random forests, which can more naturally accommodate nonlinear structures and intricate dependencies among omics variables.

While all three IMD-based methods performed well, there were subtle differences in their behavior. The IMD-filter method tended to be slightly more conservative, often resulting in a smaller set of selected variables. This can be useful when one wishes to prioritize specificity over sensitivity. The IMD-mixture approach struck a good balance for many scenarios, effectively distinguishing signal from noise without imposing strict thresholds. Meanwhile, IMD-transformation provided a distributional normalization that helped reveal strong variables even when mean IMD values were low or tightly clustered. Researchers may choose among these methods based on their own data characteristics and objectives.

In summary, our simulation studies confirmed that the MRF-IMD framework can identify cross-correlated variables reliably under a wide array of conditions, outperforming several well-established methods. These results suggest that by embracing nonlinear modeling, leveraging IMD-based variable selection, and providing flexible selection strategies, our framework is well-suited for integrative multi-omics analysis. This strong performance in controlled simulations sets the stage for more complex real-world applications and further supports the potential utility of MRF-IMD methods in guiding biomarker discovery and driving meaningful biological interpretations.

### Individual Cancer Data: Breast Cancer and Colorectal Cancer

After selecting the optimal connections between omics datasets, we applied our MRF-based framework to breast invasive carcinoma (BRCA) and colon adenocarcinoma (COAD) data from TCGA. For BRCA, we examined three directional models: *gene* **←** *irn, et* **←** *irn*, and *irn* **←** *et*, and for COAD we considered *gene* **←** *irn, et* **←** *irn*, and *irn* **←** *gene* configuration. To assess the stability of the identified variables, we repeated the analysis 10 times for each combination and applied the three IMD-based selection methods (filter, mixture, and transformation).

Box plots in Supplementary Figure S7 show the number of selected variables across these 10 replicates. All three methods consistently produced a reasonable set of variables for both BRCA and COAD. Among the IMD-based strategies, the IMD-transformation approach tended to be the most stable, while the IMD-filter and IMD-mixture methods often selected fewer variables, illustrating a trade-off between stability and stringency.

Figure 5a displays the top 20 variables identified by the IMD-filter method in BRCA. Prominent genes included *SAMD5, MMP11*, and *GABRE. MMP11* is well-documented in the literature as playing a pivotal role in breast cancer, showing high expression levels in early luminal subtypes^27,28^. Additionally, *GABRE* has also been shown to be related to triple-negative breast cancer^29^. Furthermore, *ESR1*, the gene that encodes the estrogen receptor (ER) along with pioneering transcription factors *GATA3* and *FOXA1* are well established factors in hormonally dependent breast cancer^30^.

**Figure 5.**
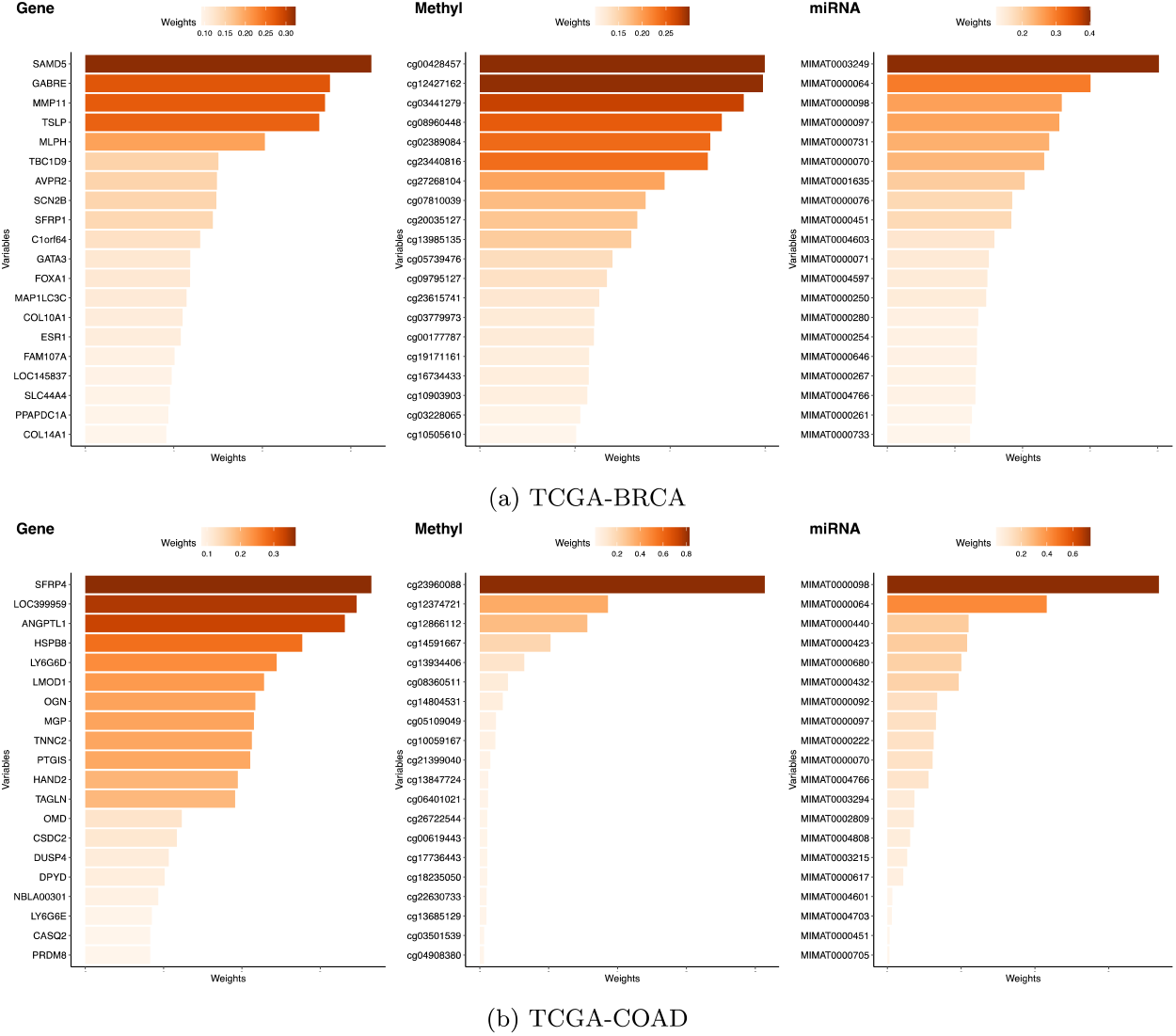
Top 20 variable weights chosen by one IMD-filter model. Many of these variables are significantly associated with (a) TCGA-BRCA and (b) TCGA-COAD.

Among the DNA methylation features, probes such as cg03441279 in *BCL9* and cg12427162 in *SFT2D2* have been associated with breast cancer prognosis^31^. On the miRNA side, MIMAT0003249 (hsa-miR-584-5p) and MIMAT0000064 (hsa-let-7c-5p) stood out, both having been previously implicated in breast cancer biology^32,33^.

We then applied the functional enrichment analysis to the selected genes with one IMD-filter model. The functional analysis was performed using *clusterProfiler* R package focusing on Canonical (C2:CP), GO (C5:BP) and Hallmark pathways from the Molecular Signatures Database^30^. Figure 6 displays the top 10 significant pathways (i.e. FDR < 0.25). Among these, several pathways are related to the estrogen receptors, which are significantly enriched and closely associated with breast cancer.

**Figure 6.**
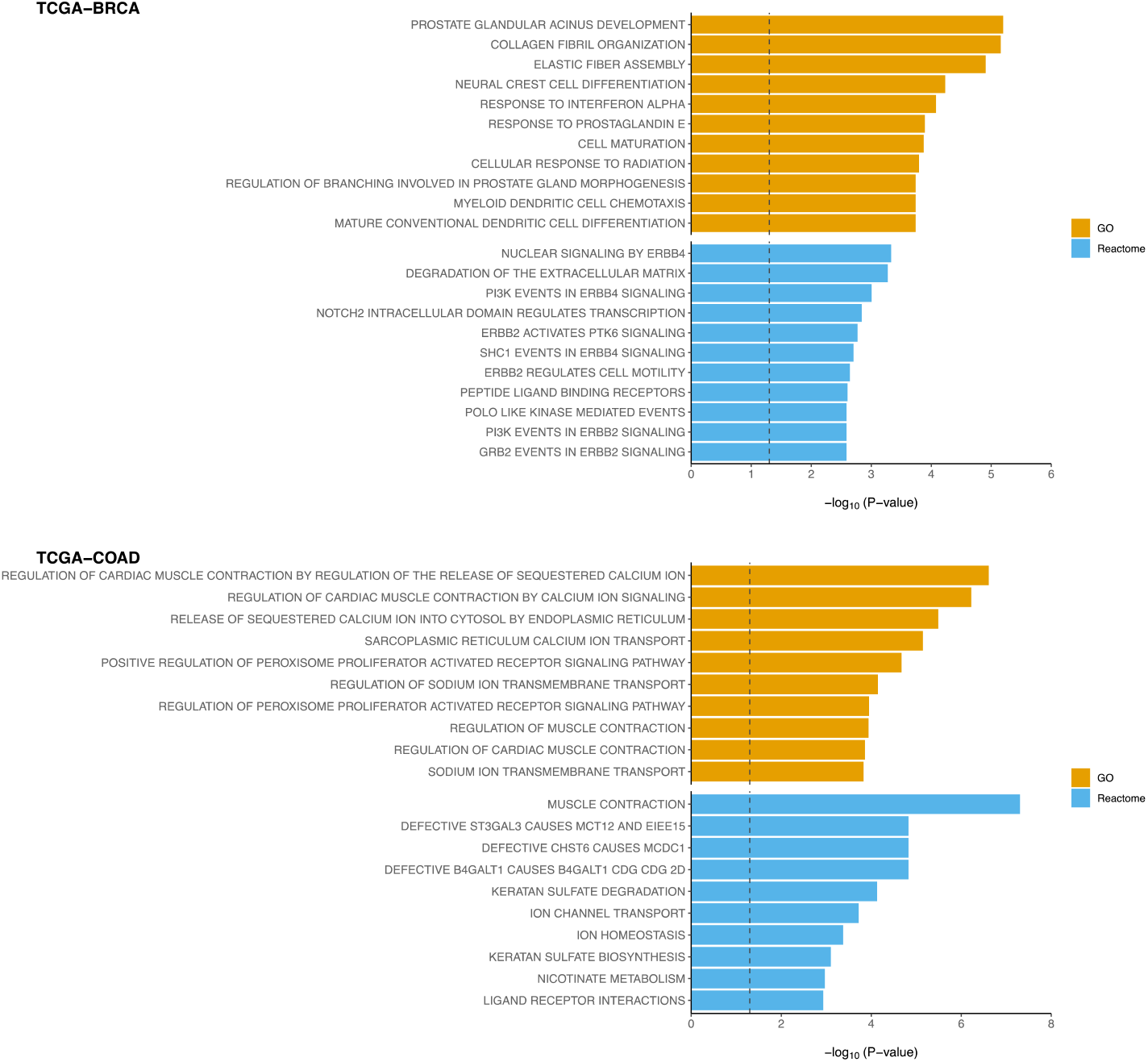
Functional enrichment analysis to the selected genes with one IMD-filter model performed using clusterProfiler R package. Top 10 significant pathways of C2:CP, GO and Hallmark (i.e. FDR < 0.25) were displayed for both TCGA-BRCA and TCGA-COAD.

For the TCGA-COAD dataset (Figure 5b), two of the top genes selected by the model—*SFRP4* and *ANGPTL1*—are known to be highly expressed in colorectal cancer (CRC) and have been linked to poor clinical outcomes in CRC patients^35–37^. Although fewer DNA methylation probes emerged prominently, cg23960088 exceeded the 0.5 weight threshold. For miRNAs, MIMAT0000098 and MIMAT0000064 were the top features selected. Functional enrichment analysis (Figure 6, bottom) revealed pathways related to muscle contraction and ion homeostasis, processes that have been connected to colorectal cancer progression.^38^.

An important goal in cancer multi-omics studies is to identify biomarkers that not only reflect biological mechanisms but also correlate with clinical outcomes. To evaluate the prognostic value of the variables identified by our MRF-based methods, we applied integrative non-negative matrix factorization (IntNMF) method^39^ to combine the selected variables from the three omics data types for both BRCA and COAD. This integration allowed us to cluster patients into two groups representing high- and low-risk survival profiles.

The Kaplan–Meier survival curves (Figure 7) for these patient clusters showed that all three MRF-IMD methods produced feature sets that robustly distinguished patient subgroups with significantly different survival rates. The log-rank tests were highly significant, suggesting that our selected biomarkers not only have biological relevance but may also hold promise for clinical risk stratification.

**Figure 7.**
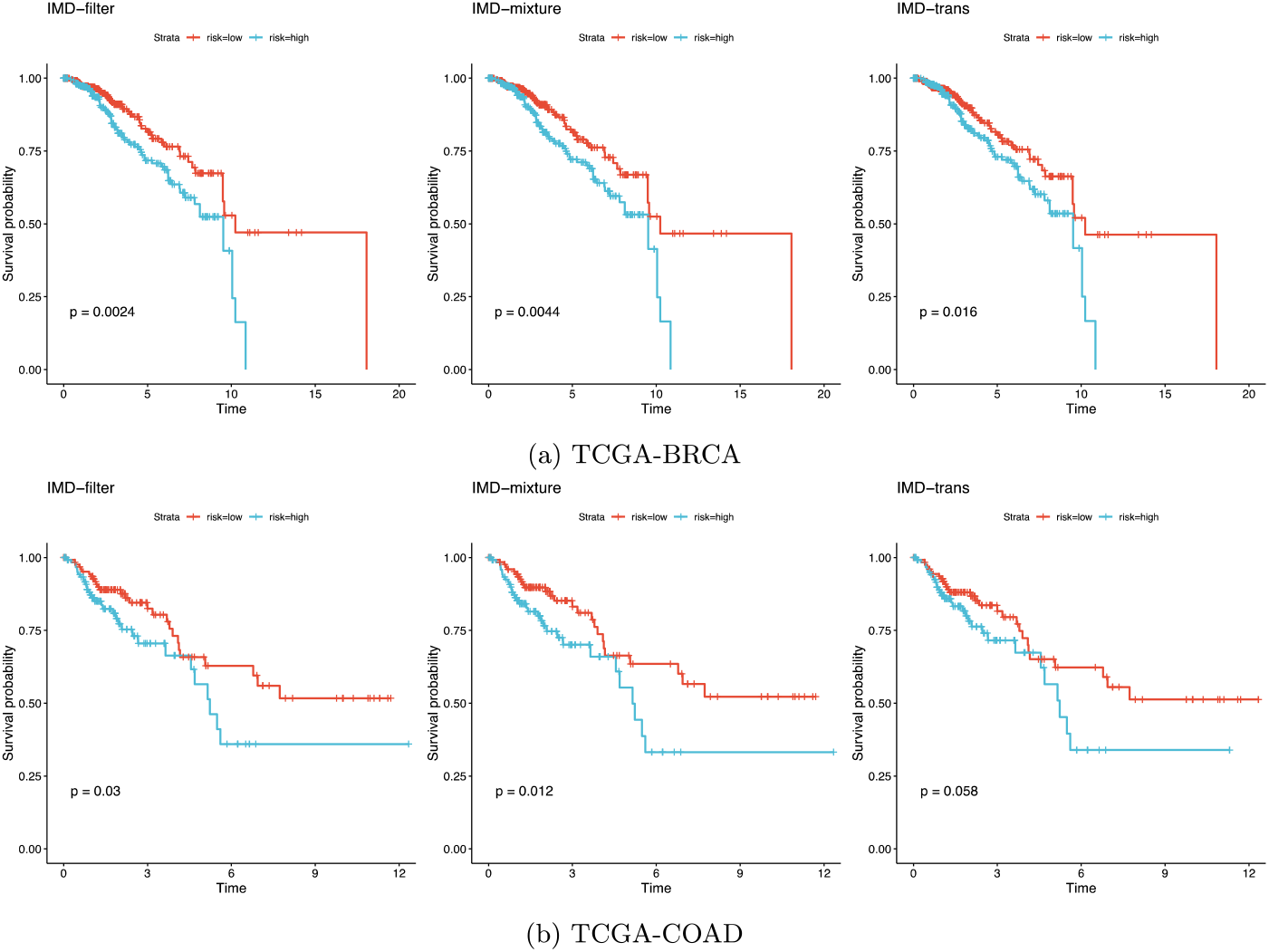
Kaplan-Meier plots and log-rank p-values for the survival analysis of clustered groups using selected integrated features. We found that all the clustering methods are robust with different variable selection methods, resulting in strongly significant log-rank p-values in both (a) TCGA-BRCA and (b) TCGA-COAD.

In summary, applying our MRF-based framework to BRCA and COAD datasets uncovered key genes, methylation probes, and miRNAs that are consistent with known cancer biology. The enriched pathways and strong associations with survival outcomes highlight the potential of these methods to identify meaningful biomarkers in multi-omics data, guiding future research and, potentially, clinical translation.

### TCGA PAN Cancer Data

To further demonstrate the flexibility and scalability of our framework, we applied the IMD-transformation variable selection method to a pan-cancer dataset from TCGA, encompassing 22 distinct cancer types. After processing and feature selection, we retained 186 ATAC-seq features and 300 RNA-seq features for integrative analysis. We first visualized the data using Uniform Manifold Approximation and Projection (UMAP), which revealed clear and distinct clusters that corresponded to different cancer types (Figure 8).

**Figure 8.**
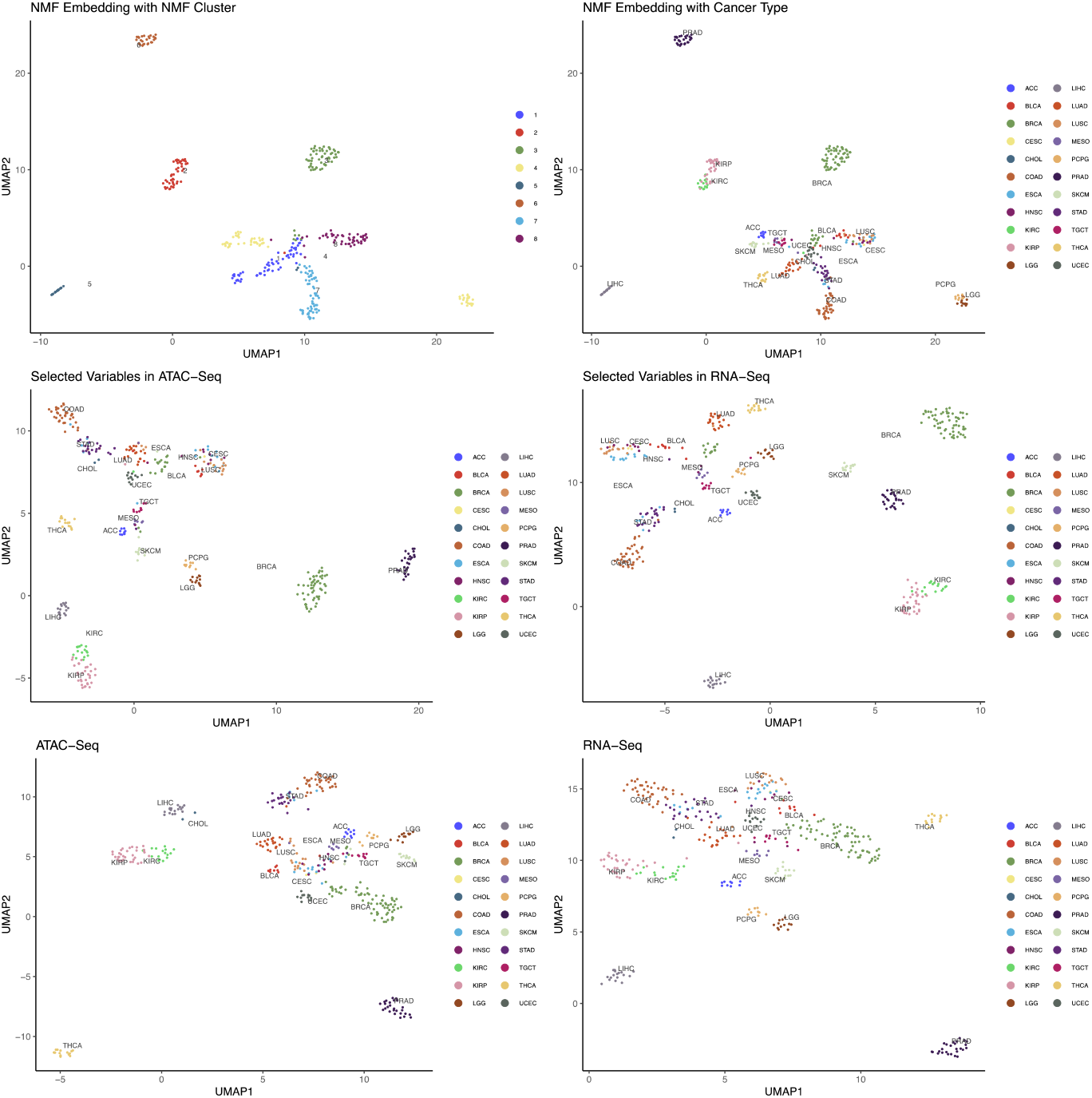
Two-dimensional UMAP embedding of TCGA PAN cancer data. (a) and (b) are UMAP of NMF embeddings of 300 RNA-Seq and 186 ATAC-Seq features and colored by NMF clustering groups and 22 PAN cancer types. (c) and (d) are UMAP of selected RNA-Seq and ATAC-Seq features colored by the 22 PAN cancer types. (e) and (f) are UMAP of 5,000 RNA-Seq and 50,000 ATAC-Seq features. Here we can see that the selected features have already captured the distinction between different cancer types.

Next, we employed integrative non-negative matrix factorization (IntNMF) to identify biologically meaningful groups within this large, heterogeneous dataset. We determined the optimal number of clusters using the nmf.opt.k function from the IntNMF R package. Figure 9 shows the resulting confusion matrix, illustrating how our selected features effectively separated the samples into eight clusters. Each cluster highlighted unique molecular characteristics and captured established patterns of tumor heterogeneity. For example:

**Figure 9.**
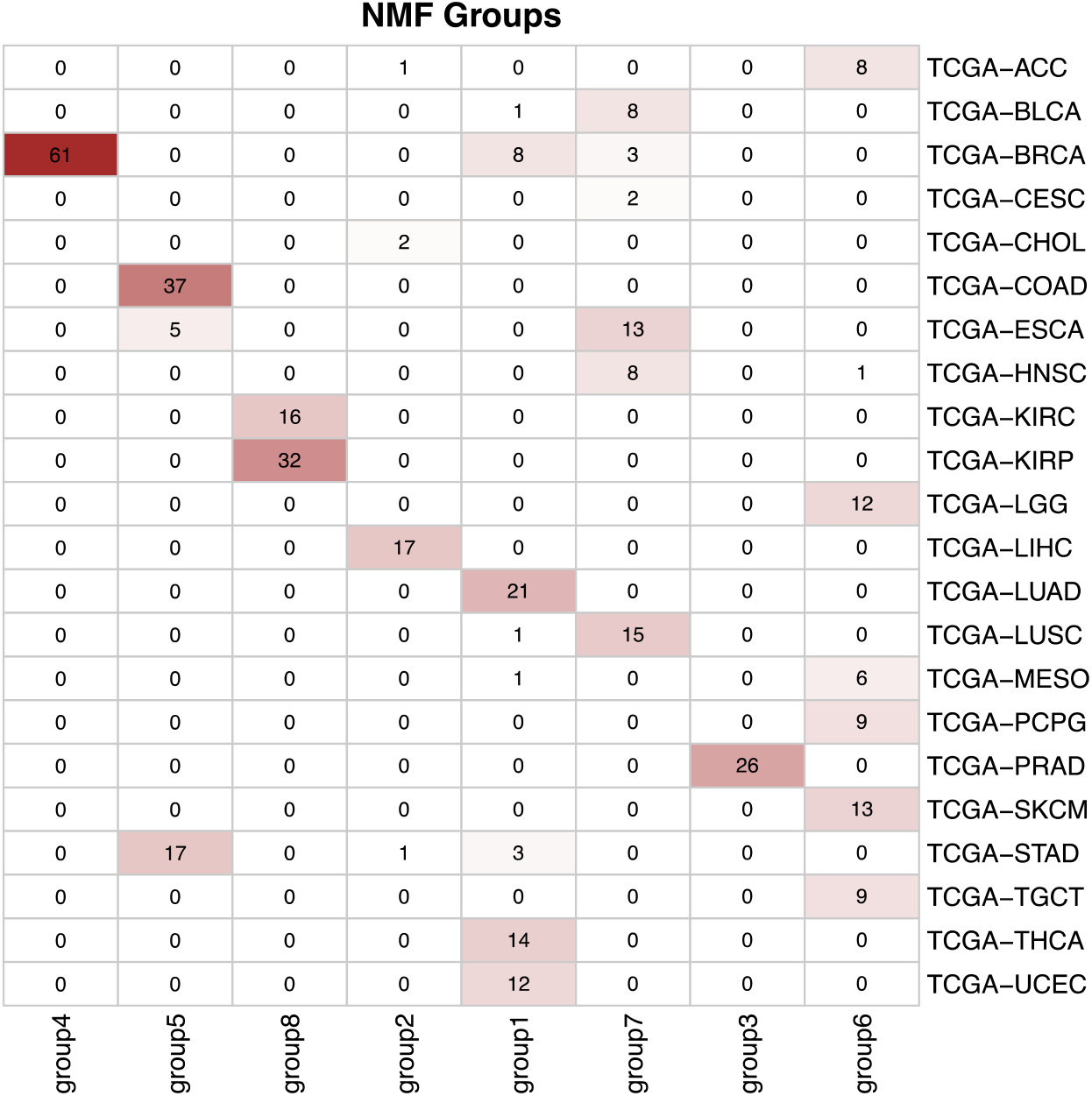
Confusion matrix of IntNMF clustering results of selected ATAC-seq and RNA-seq. The variable selection process effectively identified key features across the multi-omics datasets, distinguishing between the cancer types.

- **Group 1 (BBRCA-UCEC Cluster)**: This cluster, which includes basal-like breast tumors and uterine corpus endometrial carcinoma (UCEC), is marked by high genomic instability and frequent TP53 and BRCA1 mutations. These molecular signatures underscore the importance of DNA repair and cell cycle regulation pathways.
- **Group 4 (Non-basal Breast Cancer Cluster)**: Enriched for luminal and HER2-positive breast cancers, this group points to differences in hormone receptor signaling and distinct oncogenic pathways, aligning with known breast cancer subtypes and their varying treatment responses.
- **Group 5 (Gastrointestinal Adenocarcinoma CIN, GA-CIN)**: Consisting of colon, stomach, and esophageal adenocarcinomas, this cluster features tumors originating from the epithelial lining of the gastrointestinal tract. They share chromosomal instability (CIN) and frequent genetic alterations, reflecting well-known drivers of gastrointestinal cancer progression.
- **Group 7 (Squamous Cell Cluster, SCC)**: This group unites several squamous cell carcinomas (e.g., esophageal, head and neck, lung, and bladder). Common alterations in TP53, combined with genomic instability and activation of RTK-RAS and PI3K pathways, highlight conserved mechanisms underlying squamous cell tumorigenesis.
- **Group 8 (Renal Epithelial Carcinomas, REC)**: Kidney renal papillary cell carcinoma (KIRP) and kidney renal clear cell carcinoma (KIRC) formed a distinct cluster characterized by features inherent to the renal epithelium. This pattern reflects their shared origin and common molecular hallmarks.

By capturing these known and biologically meaningful patterns, our MRF-IMD framework shows its utility in differentiating tumor types, identifying key molecular signatures, and offering a more integrated view of tumor diversity. These results illustrate the method’s promise for guiding future studies on cancer classification, patient stratification, and uncovering novel therapeutic targets across a wide range of malignancies.

## DISCUSSION

The continuous growth of high-throughput technologies has enabled the profiling of multiple omics layers—spanning the genome, epigenome, transcriptome, and beyond—within the same samples. This multi-omics landscape offers the potential for more complete insights into disease mechanisms, biomarker discovery, and therapeutic targeting. However, integrating these disparate data types and identifying meaningful shared features remains an ongoing challenge.

In this study, we presented a multivariate random forest (MRF)-based framework enhanced by the inverse minimal depth (IMD) metric to address these challenges. By combining the strengths of MRF for capturing nonlinear relationships and the IMD-based strategies for feature selection, our approach provides a flexible and robust solution for multi-omics integration. Unlike conventional linear methods such as sPLS and CCA, which often assume simpler data structures and can be prone to overfitting, the MRF-IMD framework scales well to complex, high-dimensional scenarios.

Our simulation studies showed that MRF-IMD methods consistently outperform established approaches in identifying cross-correlated variables. The results held true under diverse conditions, including both linear and nonlinear models, varying sample sizes, and multiple levels of dimensionality and noise. These findings highlight the robustness and adaptability of our method, reinforcing its suitability for real-world applications.

We further demonstrated the framework’s utility using multi-omics data from The Cancer Genome Atlas (TCGA). In breast and colorectal cancer, our approach uncovered known cancer-related genes, miRNAs, and DNA methylation features, as well as biologically relevant pathways. Moreover, clustering patients based on selected variables revealed groups with distinct survival outcomes, underscoring the clinical relevance of our discoveries. In a pan-cancer setting, we showed that the MRF-IMD method could detect key molecular differences among diverse tumor types, identifying clusters with characteristic genomic instabilities, pathway alterations, and tissue-of-origin patterns. These results suggest that the approach has broad applicability, enhancing our understanding of tumor heterogeneity and potential therapeutic targets.

While our method provides clear advantages, some limitations remain. Computation time may increase with more datasets and extreme high-dimensionality. Future research could focus on improving efficiency, potentially through parallelization or dimensionality reduction strategies that preserve essential biological signals. Additionally, further integration with downstream validation steps—such as experimental verification or functional assays—would help confirm the biological significance of the selected variables and strengthen the evidence for potential biomarkers.

In conclusion, our MRF-IMD framework represents a step forward in integrative multi-omics analysis. By balancing flexibility, scalability, interpretability, and robustness, it enables researchers to identify meaningful biomarkers and pathways that would be difficult to pinpoint using conventional approaches. As the field continues to generate increasingly complex data, methods like ours will be instrumental in translating multi-omics information into actionable insights that advance our understanding of health and disease.

## Supporting information

SuppFigs

SuppNotes

## DATA AVAILABILITY

- TCGA-BRCA
  - mRNA expression data (Gene): https://xenabrowser.net/datapages/?dataset=TCGA.BRCA.sampleMap%2FHiSeqV2&host=https%3A%2F%2Ftcga.xenahubs.net&removeHub=https%3A%2F%2Fxena.treehouse.gi.ucsc.edu%3A443
  - miRNA expression data (miRNA): https://xenabrowser.net/datapages/?dataset=TCGA.BRCA.sampleMap%2FmiRNA_HiSeq_gene&host=https%3A%2F%2Ftcga.xenahubs.net&removeHub=https%3A%2F%2Fxena.treehouse.gi.ucsc.edu%3A443
  - DNA methylation data (Methyl): https://xenabrowser.net/datapages/?dataset=TCGA.BRCA.sampleMap%2FHumanMethylation450&host=https%3A%2F%2Ftcga.xenahubs.net&removeHub=https%3A%2F%2Fxena.treehouse.gi.ucsc.edu%3A443
- TCGA-COAD
  - mRNA expression data (Gene): https://xenabrowser.net/datapages/?dataset=TCGA.COAD.sampleMap%2FHiSeqV2&host=https%3A%2F%2Ftcga.xenahubs.net&removeHub=https%3A%2F%2Fxena.treehouse.gi.ucsc.edu%3A443
  - miRNA expression data (miRNA): https://xenabrowser.net/datapages/?dataset=TCGA.COAD.sampleMap%2FmiRNA_HiSeq_gene&host=https%3A%2F%2Ftcga.xenahubs.net&removeHub=https%3A%2F%2Fxena.treehouse.gi.ucsc.edu%3A443
  - DNA methylation data (Methyl): https://xenabrowser.net/datapages/?dataset=TCGA.COAD.sampleMap%2FHumanMethylation450&host=https%3A%2F%2Ftcga.xenahubs.net&removeHub=https%3A%2F%2Fxena.treehouse.gi.ucsc.edu%3A443
- TCGA-Pan-Cancer
  - ATAC sequencing data (ATACseq): https://xenabrowser.net/datapages/?dataset=TCGA_ATAC_peak_Log2Counts_dedup_sample&host=https%3A%2F%2Fatacseq.xenahubs.net&removeHub=https%3A%2F%2Fxena.treehouse.gi.ucsc.edu%3A443
  - RNA sequencing (RNAseq) data downloading code: https://github.com/TransBioInfoLab/multiRF-vs/blob/main/code/real_data/data_prepare.Rmd
- Access:
  - RNAseq datasets: from R package TCGAbiolinks
  - Others: from UCSC Xena: https://xena.ucsc.edu/

## AVAILABILITY OF SUPPORTING SOURCE CODE AND REQUIREMENTS

Project name: An Integrative Multi-Omics Random Forest Framework for Robust Biomarker Discovery

Project homepage: https://github.com/TransBioInfoLab/multiRF-vs

Vignette: https://rpubs.com/noblegasss/multiRF-vs-vignette

Operating system: Linux (Ubuntu)

Programming language: R version 4.4.2

License: GPL 3.0 or higher

## SUPPLEMENTARY DATA

Supplementary Data are available at GigaScience online.

## REFERENCES

1 Singh, A., Shannon, C.P., Gautier, B., Rohart, F., Vacher, M., Tebbutt, S.J., and Lê Cao, K.-A. (2019). DIABLO: an integrative approach for identifying key molecular drivers from multi-omics assays. Bioinformatics 35, 3055–3062. 10.1093/bioinformatics/bty1054.

2 Wang, T., Shao, W., Huang, Z., Tang, H., Zhang, J., Ding, Z., and Huang, K. (2021). MOGONET integrates multi-omics data using graph convolutional networks allowing patient classification and biomarker identification. Nat Commun 12, 3445. 10.1038/s41467-021-23774-w.

3 Xiao, L., Zhang, F., and Zhao, F. (2022). Large-scale microbiome data integration enables robust biomarker identification. Nat Comput Sci 2, 307–316. 10.1038/s43588-022-00247-8.

4 Coletti, R., and Lopes, M.B. (2023). Multi-omics Data Integration and Network Inference for Biomarker Discovery in Glioma. In Progress in Artificial Intelligence, N. Moniz, Z. Vale, J. Cascalho, C. Silva, and R. Sebastião, eds. (Springer Nature Switzerland), pp. 247–259. 10.1007/978-3-031-49011-8_20.

5 Wold, H. (1966). Estimation of principal components and related models by iterative least squares.

6 Chun, H., and Keleş, S. (2010). Sparse partial least squares regression for simultaneous dimension reduction and variable selection. Journal of the Royal Statistical Society: Series B (Statistical Methodology) 72, 3–25. 10.1111/j.1467-9868.2009.00723.x.

7 Hotelling, H. (1936). Relations Between Two Sets of Variates. Biometrika 28, 321–377. 10.2307/2333955.

8 Witten, D.M., Tibshirani, R., and Hastie, T. (2009). A penalized matrix decomposition, with applications to sparse principal components and canonical correlation analysis. Biostatistics 10, 515–534. 10.1093/biostatistics/kxp008.

9 Tenenhaus, A., and Tenenhaus, M. (2011). Regularized Generalized Canonical Correlation Analysis. Psychometrika 76, 257–284. 10.1007/s11336-011-9206-8.

10 Lai, P.L., and Fyfe, C. (2000). Kernel and nonlinear canonical correlation analysis. Int. J. Neur. Syst. 10, 365–377. 10.1142/S012906570000034X.

11 Yoshida, K., Yoshimoto, J., and Doya, K. (2017). Sparse kernel canonical correlation analysis for discovery of nonlinear interactions in high-dimensional data. BMC Bioinformatics 18, 108. 10.1186/s12859-017-1543-x.

12 Breiman, L. (2001). Random Forests. Machine Learning 45, 5–32. 10.1023/A:1010933404324.

13 Segal, M., and Xiao, Y. (2011). Multivariate random forests. WIREs Data Mining and Knowledge Discovery 1, 80–87. 10.1002/widm.12.

14 Gregorutti, B., Michel, B., and Saint-Pierre, P. (2017). Correlation and variable importance in random forests. Stat Comput 27, 659–678. 10.1007/s11222-016-9646-1.

15 Diaz-Uriarte, R. (2007). GeneSrF and varSelRF: a web-based tool and R package for gene selection and classification using random forest. BMC Bioinformatics 8, 328. 10.1186/1471-2105-8-328.

16 Ishwaran, H., Kogalur, U.B., Blackstone, E.H., and Lauer, M.S. (2008). Random survival forests. Ann. Appl. Stat. 2. 10.1214/08-AOAS169.

17 Ishwaran, H., Kogalur, U.B., Gorodeski, E.Z., Minn, A.J., and Lauer, M.S. (2010). High-Dimensional Variable Selection for Survival Data. Journal of the American Statistical Association 105, 205–217. 10.1198/jasa.2009.tm08622.

18 Tang, F., and Ishwaran, H. (2017). Random Forest Missing Data Algorithms. Stat Anal Data Min 10, 363–377. 10.1002/sam.11348.

19 Ishwaran, H., Kogalur, U.B., Chen, X., and Minn, A.J. (2011). Random survival forests for high-dimensional data. Statistical Analysis and Data Mining: The ASA Data Science Journal 4, 115–132. 10.1002/sam.10103.

20 Dempster, A.P., Laird, N.M., and Rubin, D.B. (1977). Maximum Likelihood from Incomplete Data Via the EM Algorithm. Journal of the Royal Statistical Society: Series B (Methodological) 39, 1–22. 10.1111/j.2517-6161.1977.tb01600.x.

21 Lee, G., and Scott, C. (2012). EM algorithms for multivariate Gaussian mixture models with truncated and censored data. Computational Statistics & Data Analysis 56, 2816–2829. 10.1016/j.csda.2012.03.003.

22 Degenhardt, F., Seifert, S., and Szymczak, S. (2019). Evaluation of variable selection methods for random forests and omics data sets. Briefings in Bioinformatics 20, 492–503. 10.1093/bib/bbx124.

23 Cancer Genome Atlas Network (2012). Comprehensive molecular portraits of human breast tumours. Nature 490, 61–70. 10.1038/nature11412.

24 Cancer Genome Atlas Network (2012). Comprehensive molecular characterization of human colon and rectal cancer. Nature 487, 330–337. 10.1038/nature11252.

25 Corces, M.R., Granja, J.M., Shams, S., Louie, B.H., Seoane, J.A., Zhou, W., Silva, T.C., Groeneveld, C., Wong, C.K., Cho, S.W., et al. (2018). The chromatin accessibility landscape of primary human cancers. Science 362, eaav1898. 10.1126/science.aav1898.

26 Goldman, M.J., Craft, B., Hastie, M., Repecka, K., McDade, F., Kamath, A., Banerjee, A., Luo, Y., Rogers, D., Brooks, A.N., et al. (2020). Visualizing and interpreting cancer genomics data via the Xena platform. Nat Biotechnol 38, 675–678. 10.1038/s41587-020-0546-8.

27 Molière, S., Lodi, M., Leblanc, S., Gressel, A., Mathelin, C., Alpy, F., Chenard, M.-P., and Tomasetto, C. (2024). MMP-11 expression in early luminal breast cancer: associations with clinical, MRI, pathological characteristics, and disease-free survival. BMC Cancer 24, 295. 10.1186/s12885-024-11998-0.

28 Zhuang, Y., Li, X., Zhan, P., Pi, G., and Wen, G. (2021). MMP11 promotes the proliferation and progression of breast cancer through stabilizing Smad2 protein. Oncol Rep 45, 16. 10.3892/or.2021.7967.

29 Li, X., Wang, H., Yang, X., Wang, X., Zhao, L., Zou, L., Yang, Q., Hou, Z., Tan, J., Zhang, H., et al. (2021). GABRP sustains the stemness of triple-negative breast cancer cells through EGFR signaling. Cancer Lett 514, 90–102. 10.1016/j.canlet.2021.04.028.

30 Martin, E.M., Orlando, K.A., Yokobori, K., and Wade, P.A. (2021). The estrogen receptor/GATA3/FOXA1 transcriptional network: lessons learned from breast cancer. Curr Opin Struct Biol 71, 65–70. 10.1016/j.sbi.2021.05.015.

31 Xu, J., Xiang, L., Liu, Q., Gilmore, H., Wu, J., Tang, J., and Madabhushi, A. (2016). Stacked Sparse Autoencoder (SSAE) for Nuclei Detection on Breast Cancer Histopathology Images. IEEE Transactions on Medical Imaging 35, 119–130. 10.1109/TMI.2015.2458702.

32 Denkiewicz, M., Saha, I., Rakshit, S., Sarkar, J.P., and Plewczynski, D. (2019). Identification of Breast Cancer Subtype Specific MicroRNAs Using Survival Analysis to Find Their Role in Transcriptomic Regulation. Front. Genet. 10. 10.3389/fgene.2019.01047.

33 Qattan, A., Intabli, H., Alkhayal, W., Eltabache, C., Tweigieri, T., and Amer, S.B. (2017). Robust expression of tumor suppressor miRNA’s let-7 and miR-195 detected in plasma of Saudi female breast cancer patients. BMC Cancer 17, 799. 10.1186/s12885-017-3776-5.

34 Liberzon, A., Subramanian, A., Pinchback, R., Thorvaldsdóttir, H., Tamayo, P., and Mesirov, J.P. (2011). Molecular signatures database (MSigDB) 3.0. Bioinformatics 27, 1739–1740. 10.1093/bioinformatics/btr260.

35 Huang, D., Yu, B., Deng, Y., Sheng, W., Peng, Z., Qin, W., and Du, X. (2010). SFRP4 was overexpressed in colorectal carcinoma. J Cancer Res Clin Oncol 136, 395–401. 10.1007/s00432-009-0669-2.

36 Nfonsam, L.E., Jandova, J., Jecius, H.C., Omesiete, P.N., and Nfonsam, V.N. (2019). SFRP4 expression correlates with epithelial mesenchymal transition-linked genes and poor overall survival in colon cancer patients. World J Gastrointest Oncol 11, 589–598. 10.4251/wjgo.v11.i8.589.

37 Chang, T.-Y., Lan, K.-C., Chiu, C.-Y., Sheu, M.-L., and Liu, S.-H. (2022). ANGPTL1 attenuates cancer migration, invasion, and stemness through regulating FOXO3a-mediated SOX2 expression in colorectal cancer. Clin Sci (Lond) 136, 657–673. 10.1042/CS20220043.

38 Baruah, V.J., Neog Bora, P., Sarmah, B., Mahanta, P., Sarmah, A., Moretti, S., Kumar, R., and Borkotokey, S. (2022). Game-theoretic link relevance indexing on genome-wide expression dataset identifies putative salient genes with potential etiological and diapeutics role in colorectal cancer. Sci Rep 12, 13409. 10.1038/s41598-022-17266-0.

39 Chalise, P., and Fridley, B.L. (2017). Integrative clustering of multi-level ‘omic data based on non-negative matrix factorization algorithm. PLoS One 12, e0176278. 10.1371/journal.pone.0176278.

