## Supplementary material for "An Integrative Multi-Omics Random Forest Framework for Robust Biomarker Discovery": SuppFigs

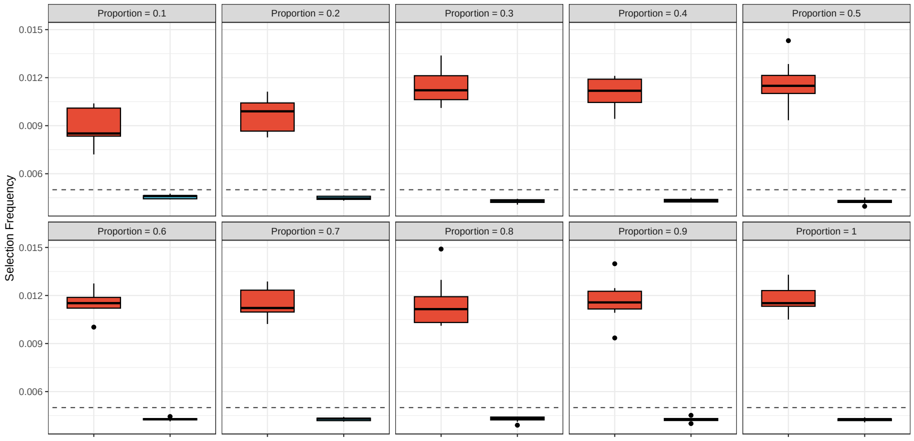


(a) Scenario 1: $q = 200$

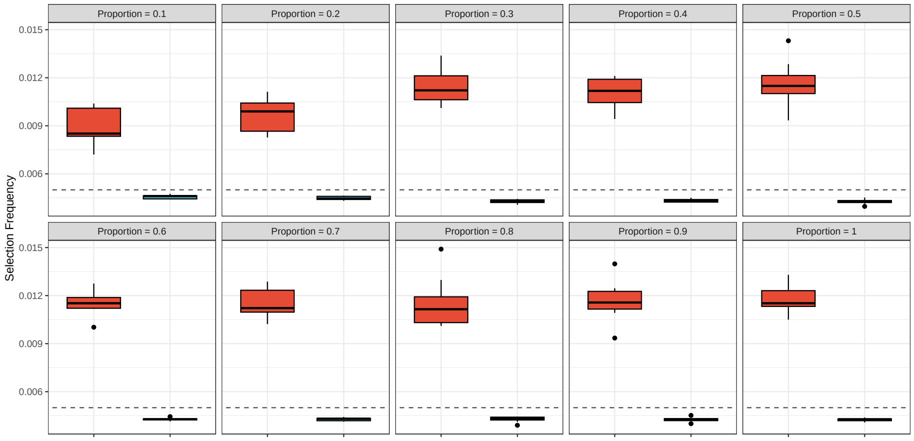


(b) Scenario 2: $q = 500$

Figure S1: Box plot of selection frequency of variables in $\mathbf{Y}$ selected as MSRV. The red box (left) represents selection frequency of cross-correlated variables and blue box (right) represents selection frequency of noise. In both scenarios, cross-correlated variables are selected as MSRV more frequently than noise variables.

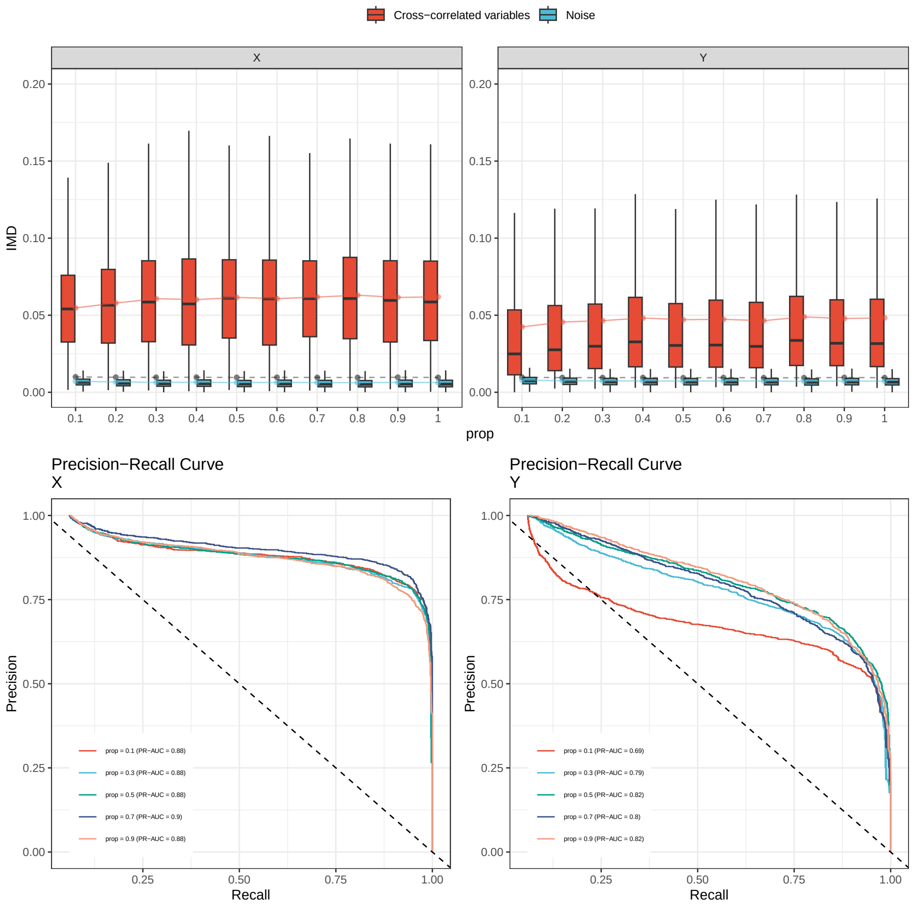


Figure S2: Impact of tree number on IMD. The top row shows box plots of IMD values for models built with varying number of trees (50, 100, 300, 500, and 1000). In both datasets, cross-correlated variables (shown in red) consistently have higher IMD values compared to noise variables (shown in blue), highlighting the greater importance of cross-correlated variables in the models. The bottom row shows the PR curve presented for each tree setting. The PR-AUC improves when ntree increases.

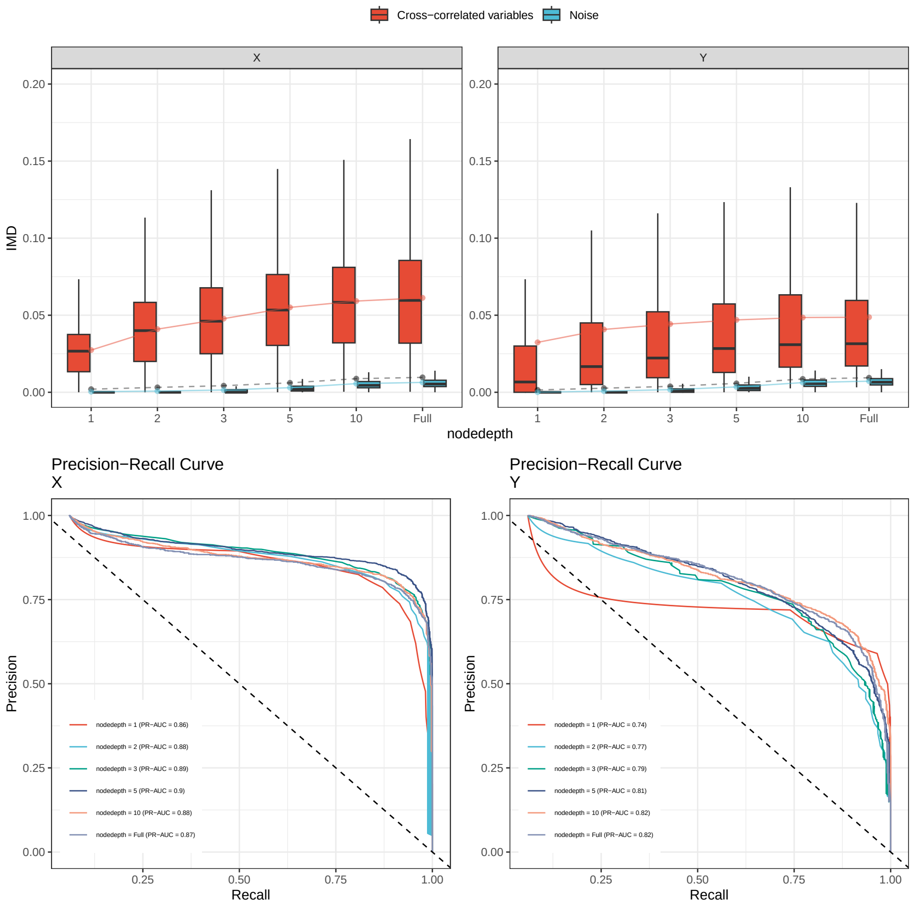


Figure S3: Impact of tree depth on IMD. The top row displays box plots of IMD values across different node depths (1, 2, 3, 5, 10, and Full). For both datasets, the mean and variation of IMD values for both cross-correlated and noise variables increase as tree depth grows. The bottom row shows the PR curve presented for each node depth setting. These results indicate that deeper trees improve variable important representation for dataset **Y**, while dataset **X** maintains high variable important representation even in very shallow trees.

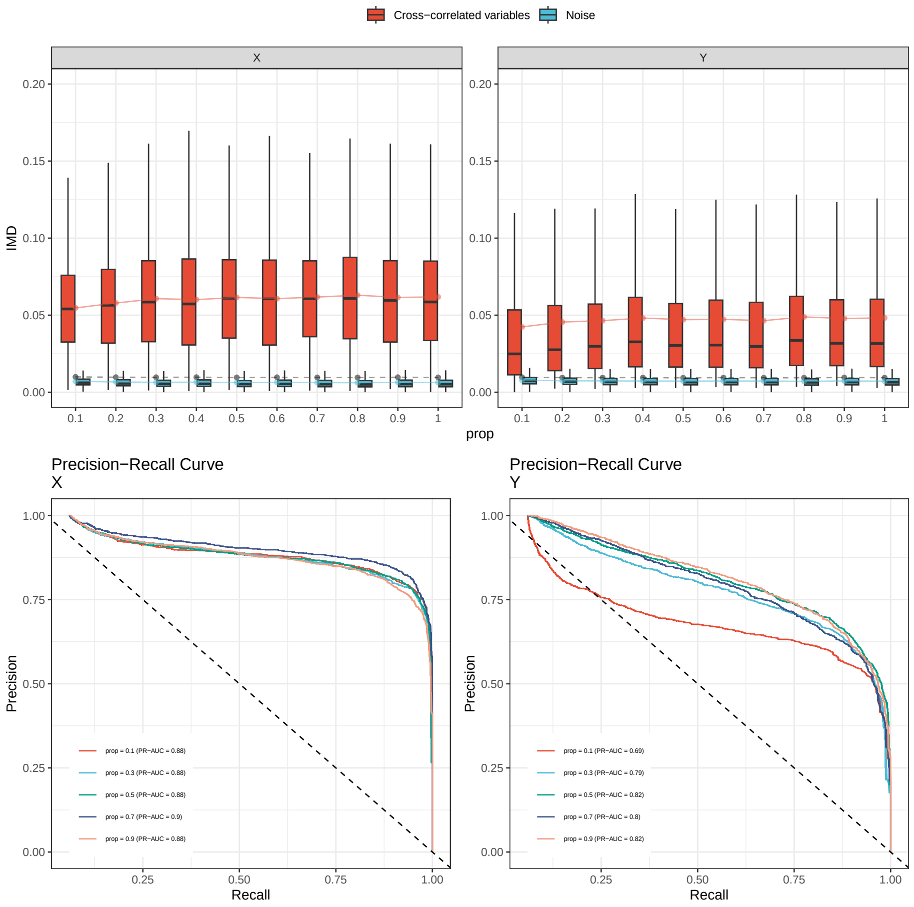


Figure S4: Impact of proportion of response variables on IMD. The top row displays box plots of IMD values across different proportion of response variables. The bottom row shows the PR curve presented for each proportion setting. The results suggest that dataset $\mathbf{Y}$ is more sensitive to the proportion of included variables, impacting its overall PR-AUC performance more significantly than dataset $\mathbf{X}$.

### Supplementary Figure 5

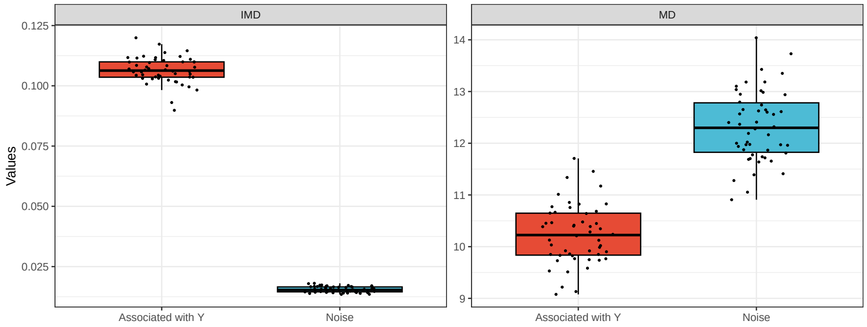


(a) Scenario 1: $q = 200$

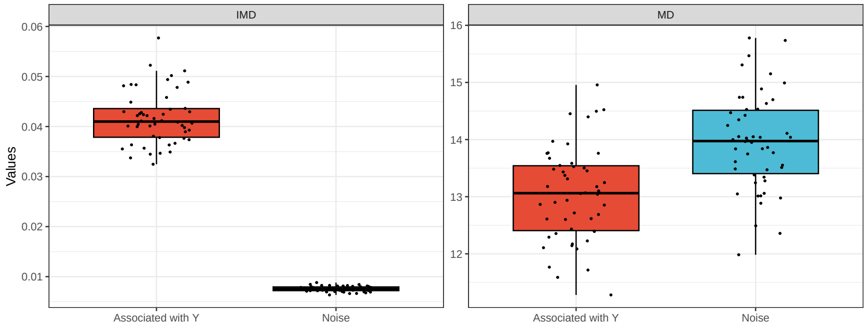


(b) Scenario 2: $q = 500$

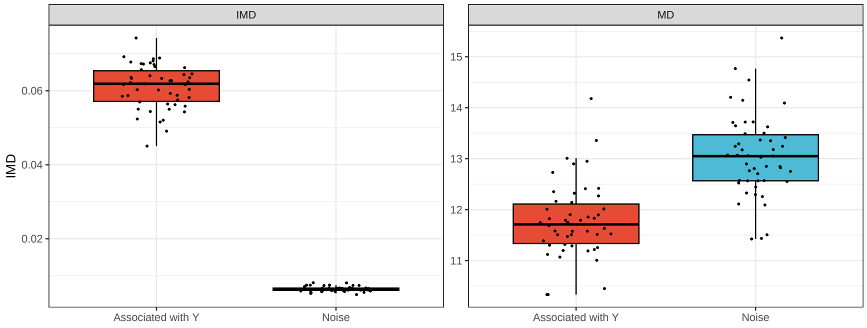


(c) Scenario 3: $q = 500$ with noise in $\boldsymbol{Y}$

Figure S5: Errorplots of the values of IMD and MD. MD shows increased variation for noise variables (in blue) across all scenarios, whereas IMD remains stable and close to zero, demonstrating consistent performance regardless of noise.

### Supplementary Figure 6

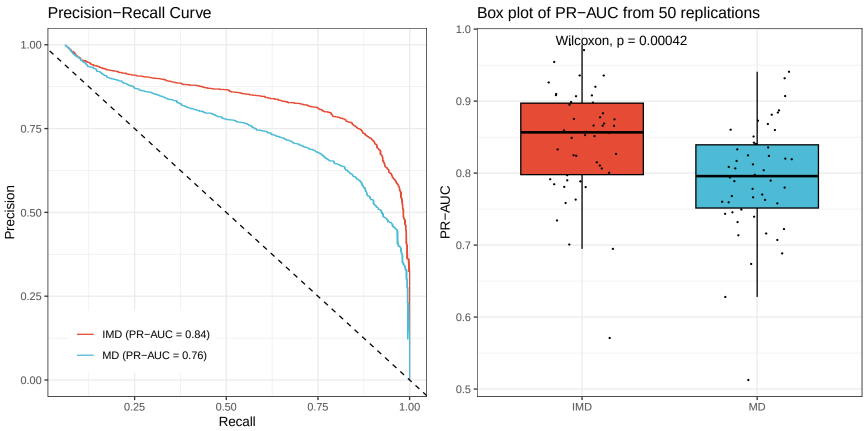


Figure S6: Compare the values of IMD and MD with the true labels of the predictors. Left: Precision-recall (PR) curve derived from 50 replications; Right: Box plot of PR-AUC obtained from each replication. The difference in performance of IMD and MD is supported by a Wilcoxon signed-rank test, which yields a significant p-value of 0.00042.

### Supplementary Figure 7

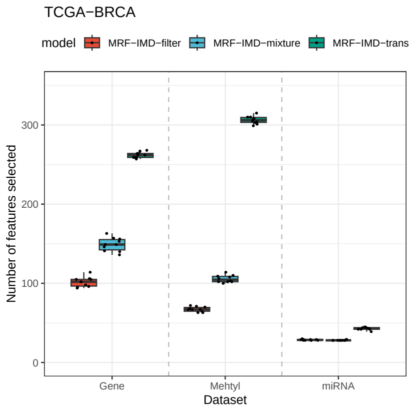


(a) TCGA-BRCA

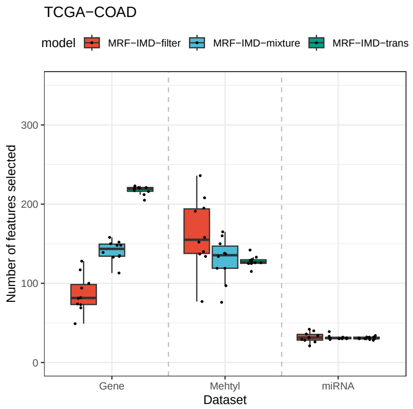


(b) TCGA-COAD

Figure S7: Boxplot of selected variables from different datasets across three methods, replicated 10 times. The results indicate that all three methods selected a reasonable number of variables across the two TCGA datasets.
