## Supplementary material for "An Integrative Multi-Omics Random Forest Framework for Robust Biomarker Discovery": SuppNotes

**Supplementary Note 1** Algorithm 1

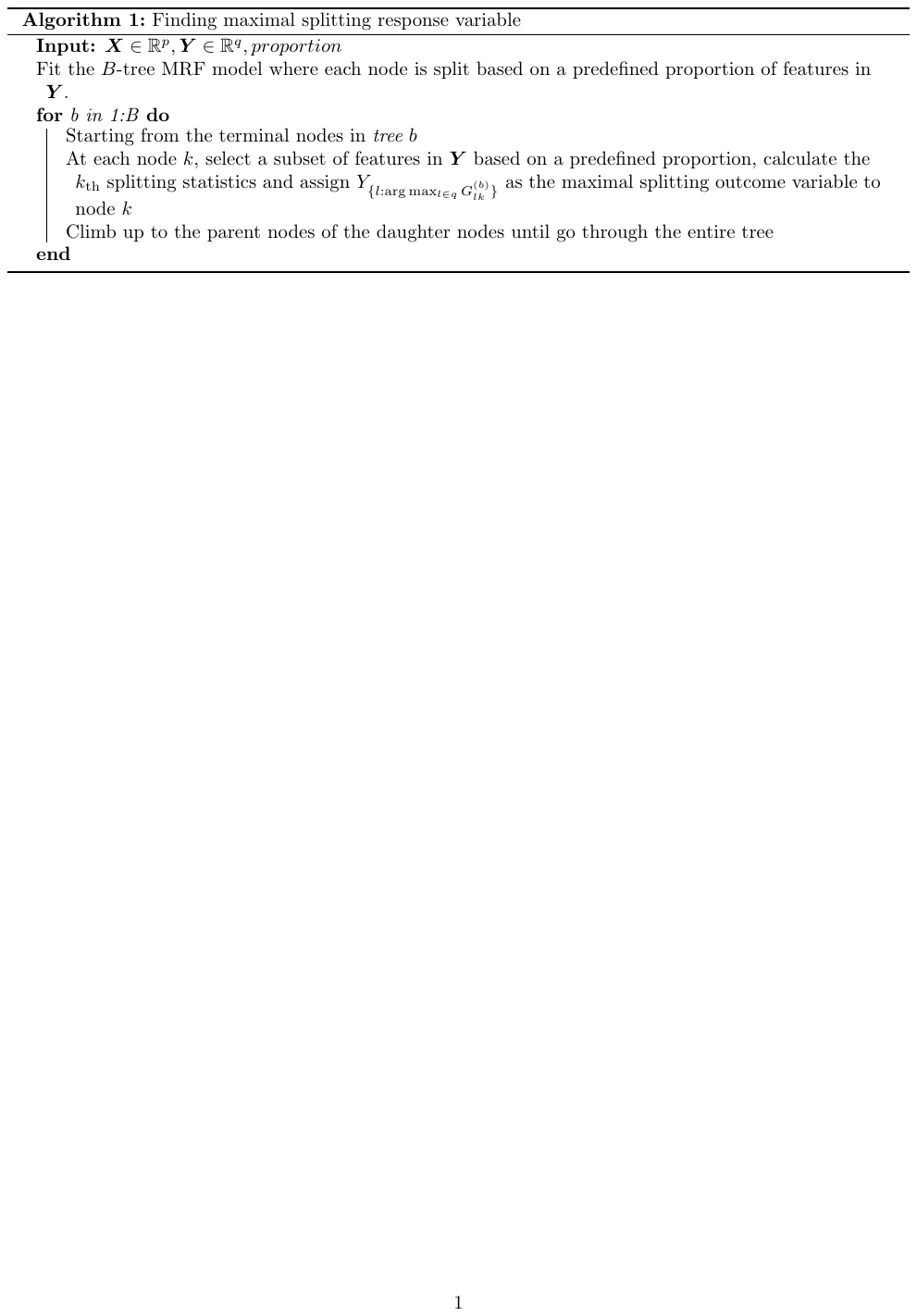

**Supplementary Note 2** Relationship between IMD And MRF Parameters

To evaluate the importance of variables in the response space, we assigned the IMD to the variables selected as the MSRV in each node. This is intuitive because the variables with lower IMD in $\mathbf{X}$ are more likely to be noise variables. As a result, when a decision tree node splits using these noise variables, it is less likely to select stronger, more influential variables from the response set $\mathbf{Y}$ as MSRV. This pattern reflects the structure of decision trees, where stronger variables, having greater predictive power, tend to appear earlier and higher in the tree. To further examine how well IMD measures variable importance across predictors and responses, we conducted exploratory analyses using various random forest parameter settings. We compared the IMD value difference between important variables and noise variables, and also evaluated model performance using Precision-Recall area under curve (PR-AUC) and PR curve under different parameter settings (see Evaluation Metrics for details). The data used in these analyses was generated from the latent model (see Simulation Study for details), $p=q=500$, $n=200$, and the first 30 variables selected as cross-correlated variables.

Supplementary Figure S2 illustrates how the number of trees (ntree) affects IMD in both datasets $\mathbf{X}$ and $\mathbf{Y}$. The top row shows box plots of IMD values for models built with varying number of trees (50, 100, 300, 500, and 1000). In both datasets, cross-correlated variables (shown in red) consistently have higher IMD values compared to noise variables (shown in blue), highlighting the greater importance of cross-correlated variables in the models. The stability of IMD values across various ntree settings indicates that the number of trees does not significantly impact the ranking of variable importance. In the bottom row, PR curves are presented for each ntree setting. For dataset $\mathbf{X}$, the PR-AUC improves as ntree increases, reaching at peak of 0.89 with 1000 trees, suggesting that model performance is enhanced with more trees. Similarly, in dataset $\mathbf{Y}$, the PR-AUC values rise with the number of trees, reaching a maximum of 0.85 at 1000 trees. These findings are presented for each ntree settings. suggest that higher ntree values improve model performance to identify important variables. However, the PR-AUC values for dataset $\mathbf{Y}$ are significantly lower when ntree $<300$, indicating that at least 300 trees should be used to accurately represent variable importance with IMD.

Supplementary Figure S3 assesses the influence of tree depth (nodedepth) on IMD values. The top row displays box plots of IMD values across different node depths (1, 2, 3, 5, 10, and Full). For both datasets, the mean and variation of IMD values for both cross-correlated and noise variables increase as tree depth grows. When nodedpeth = 1, the noise variables have an IMD of approximately 0. For dataset $\mathbf{X}$, PR-AUC values remain consistently high, ranging from 0.86 to 0.87, with little variation across different depths. In contrast, for dataset $\mathbf{Y}$, PR-AUC values increase with depth, peaking at 0.83 for node depth 10 and Full, although they remain lower than those for dataset X. These results indicate that deeper trees improve variable important representation for dataset $\mathbf{Y}$, while dataset $\mathbf{X}$ maintains high variable important representation even in very shallow trees.

As shown in Supplementary Figure S4, the IMD values for cross-correlated variables remain consistently higher than those for noise variables across both datasets. The mean IMD for cross-correlated variables hovers around 0.05, while for noise variables, it is close to zero. The PR curves indicate that the performance of the models, measured by the PR-AUC, remains relatively stable for dataset $\mathbf{X}$, with values around 0.88, showing only a slight decrease in precision at high recall levels as the proportion increases. In contrast, dataset $\mathbf{Y}$ displays more variability in PR-AUC values, ranging from 0.69 to 0.82, with noticeable differences in precision at high recall levels. This suggests that dataset $\mathbf{Y}$ is more sensitive to the proportion of included variables, impacting its overall PR-AUC performance more significantly than dataset $\mathbf{X}$. However, when the proportion of response variables selected is greater than or equal to 0.3, the IMD continued to represent the important variables in $\mathbf{Y}$ effectively.

In the box plots of Supplementary Figure S3 and Supplementary Figure S4 the mean IMD is shown for cross-correlated variables (in red), noise variables (in blue), and the entire IMD mean (in dashed grey). It is evident that the mean IMD for cross-correlated variables is significantly higher than that of noise variables across all parameter settings. Notably, in the high-dimensional noise data setting, the mean IMD is slightly elevated but still approximately equivalent to the mean IMD of the noise variables. This highlights the robustness of IMD in distinguishing between cross-correlated and noise variables, even in challenging, high-dimensional scenarios.

**Supplementary Note 3** Comparison of IMD and MD

Using the latent model as described in Simulation Study section, we computed both the inverse minimal depth (IMD) and minimal depth (MD) across the forest. We introduced noise variables in the response data to evaluate their impact on both IMD and MD calculations. Each scenario was repeated 50 times to validate the results.

As shown in Supplementary Figure S5, both IMD and MD exhibit similar distributions, with IMD essentially being the inverse of MD when variables are selected in the trees. However, MD shows increased variation for noise variables (in blue) across all scenarios, whereas IMD remains stable and close to zero, demonstrating consistent performance regardless of noise. This contrast becomes more apparent as dimensionality increases or when more noise is present in the response data. These results highlight IMD’s robustness in handling higher-dimensional and noisier data, making it a reliable measure of variable importance.

For Scenario 3, we further compared the performance of two models, IMD and MD, employing the PR-AUC as the evaluative metric. The left side of Supplementary Figure S6, presents a Precision-Recall (PR) curve, showing the trade-off between precision and recall for both models based on all 50 replications. To compute the PR-AUC for MD, we applied the negative min-max scale of MD for each replication. The PR curve for the IMD model exhibits superior performance with a PR-AUC of 0.84, while the PR-AUC for the MD model is 0.76. These results suggest that IMD reliably achieves higher precision across varying levels of recall compared to MD. On the right side of Supplementary Figure S6, a box plot illustrates the distribution of PR-AUC values from the 50 replications for both models. The box plot for IMD reveals higher median PR-AUC values and less variability than that of MD, further underscoring the superior and consistent performance of IMD. This difference in performance is further supported by a Wilcoxon signed-rank test, which yields a significant p-value of 0.00042. In conclusion, these findings clearly demonstrate the superior performance of the IMD model over the MD model in terms of PR-AUC.

**Supplementary Note 4** Algorithm 2

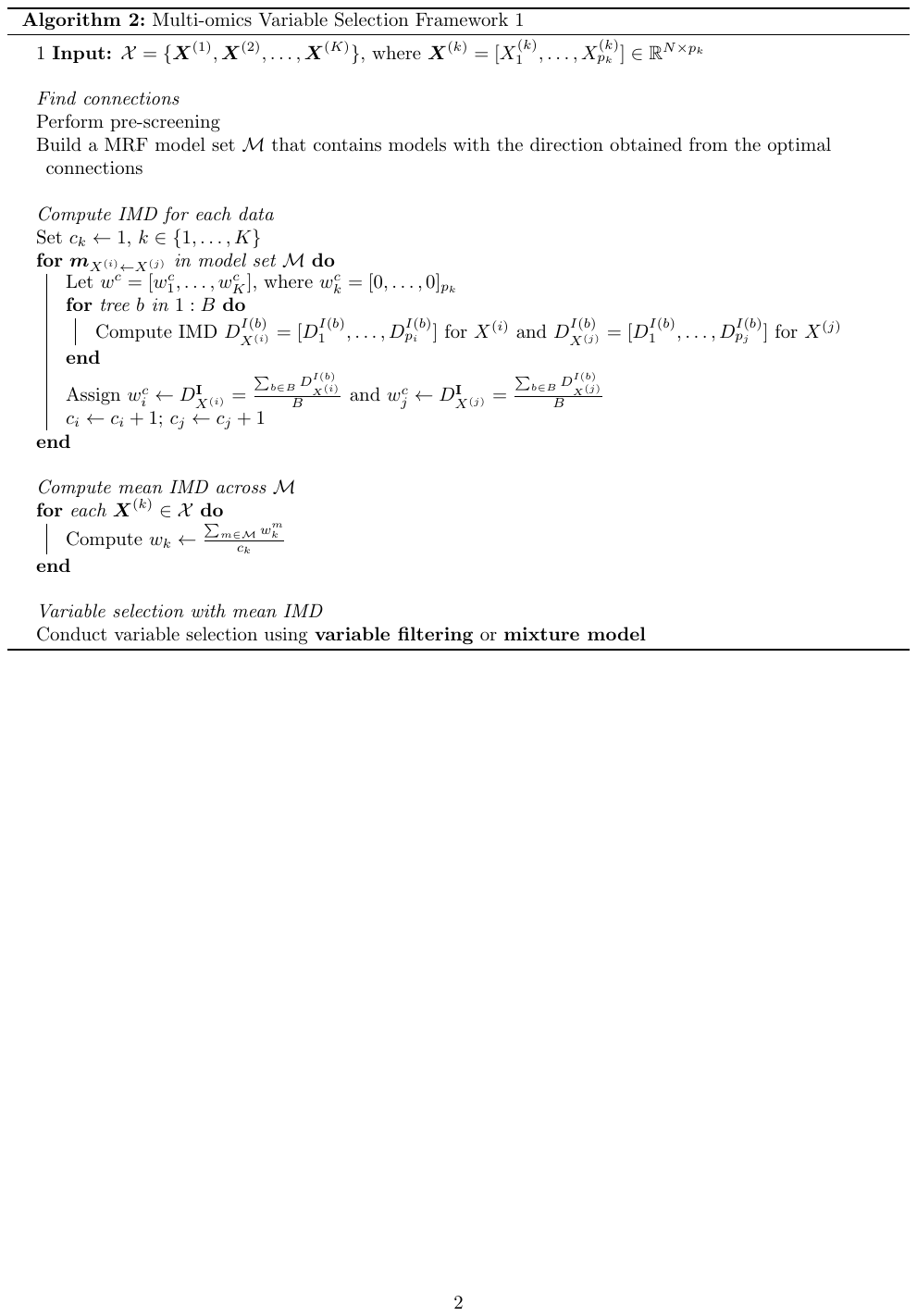

**Supplementary Note 5** Algorithm 3

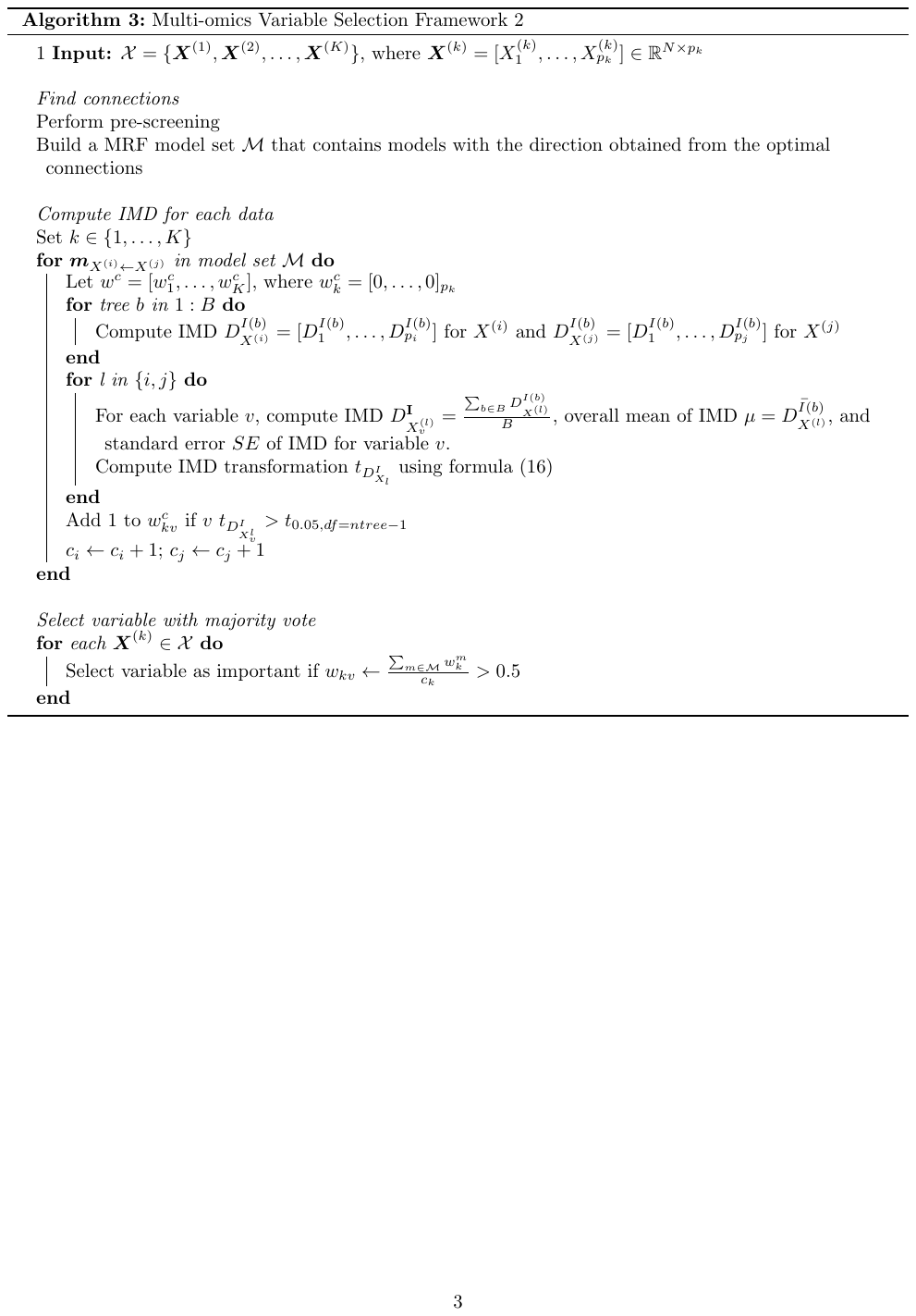
